# The relevance of GGCC methylation for cyanobacterial gene expression regulation

**DOI:** 10.64898/2026.10.20.760421

**Authors:** Nils Stappert, Nils Schmidt, Viktoria Reimann, Martin Hagemann, Wolfgang R. Hess

## Abstract

Compared to eukaryotic organisms, the regulatory significance of epigenetic DNA modifications is much less well understood in bacteria. In the cyanobacterium *Synechocystis* sp. PCC 6803, the DNA-specific methyltransferase M.Ssp6803II modifies the first cytosine in GGCC, forming N4-methylcytosine (GG^m4^CC). This motif occurs 38,512 times in the *Synechocystis* 6803 chromosome alone. Previous analyses showed that deletion of the gene encoding M.Ssp6803II (Δ*sll0729*), caused a “bluish” phenotype due to the aberrant expression of the *hemJ* gene encoding protoporphyrinogen IX oxidase, HemJ. This effect was caused by the lacking GGCC methylation in the discriminator sequence of the *hemJ* promoter in Δ*sll0729*. In the present study, we analyzed the impact of GG^m4^CC methylation on the overall gene expression. Bioinformatic analyses of multiple, independently obtained DNA microarray datasets revealed reproducible methylation-dependent changes in the expression of more than 300 genes. We selected 26 genes that had the GGCC methylation motif in different locations of their promoter sequences and verified their differential expression through qPCR analyses. Classification of the promoters according to the relative position of GGCC motifs suggested context-specific effects on gene expression. These findings imply that GG^m4^CC methylation via DNA methyltransferase M.Ssp6803II has an important impact on proper gene expression in *Synechocystis* 6803, and likely other bacteria.

## 1 Introduction

DNA base methylation occurs in all kingdoms of life. The most common types of methylated bases are endocyclic 5-methylcytosine (^m5^C), exocyclic N6-methyladenine (^m6^A), and N4-methylcytosine (^m4^C). All three methylated nucleobases have been identified in both prokaryotes and eukaryotes (Moore *et al*., 2013; Iyer *et al*., 2016; Sánchez-Romero and Casadesús, 2020; Rodriguez *et al*., 2022). While ^m5^C dominates in the eukaryotic world and is mainly localized in stretches of CpG dinucleotides (so called CpG islands), in prokaryotes ^m6^A and ^m4^C are the most abundant nucleobases to date. DNA base methylation has been found to be involved in processes as divergent as phage defense, cell cycle control, regulation of transcription, DNA replication and segregation, nucleoid organization, X-chromosome inactivation, or DNA repair and uptake (Blow *et al*., 2016; Sergeeva *et al*., 2023).

Newly methylated bases are established by DNA methyltransferases (DNMTs). These modifications do not interfere with Watson-Crick base pairing. They protrude into the major groove of the DNA double helix. This exposes the methyl group to DNA-binding proteins or the transcriptional machinery (Jeltsch, 2002). In eukaryotes, two types of DNMTs are distinguished: (i) The maintenance DNMTs which preferentially methylate hemi-methylated DNA after replication to maintain methylation patterns and (ii) the de novo DNMTs which establish new DNA methylation patterns, often within CpG islands (Dhe-Paganon *et al*., 2011; Jin and Robertson, 2013). In prokaryotes, DNMTs recognize specific DNA motifs and are commonly associated with restriction-modification (R-M) systems or act alone as single enzymes (so-called “orphan” DNMTs). Typically, DNMTs in prokaryotes target palindromic sequences and the number of different DNMTs per genome, each specific for a different recognition sequence, can vary drastically. Genes encoding DNMTs in prokaryotes were often independently acquired through horizontal gene transfer (HGT) and therefore may closely resemble each other in different taxonomic groups (Murphy *et al*., 2013). DNA methylation has been observed in more than 93% of prokaryotic genomes that were analyzed (Roberts *et al*., 2010; Blow *et al*., 2016). However, most of these DNMT systems have remained understudied with regard to their biological functions.

In both eukaryotes and prokaryotes, DNA modifications, such as ^m5^C and ^m6^A, can alter gene expression when associated with promoter sequences (Camacho and Casadesús, 2005; Militello *et al*., 2012; Kahramanoglou *et al*., 2012). While epigenetic regulation in eukaryotes is mainly mediated by CpG dinucleotides carrying a ^m5^C (Moore *et al*., 2013), other DNA methylations are at least as relevant in bacteria. For instance, the *pap*-operon of *E. coli* is regulated by ON and OFF states caused by the change in methylation of adenines in the promoter region (van der Woude *et al*., 1996). The effects of ^m4^C methylation in prokaryotes have remained less studied; Gaultney et al. (2020) demonstrated changes in transcription and pathogenicity for *Leptospira interrogans*, our previous work indicated that the missing GG^m4^CC methylation in the *hemJ* promoter affected protoporphyrinogen IX oxidase expression in the cyanobacterium *Synechocystis* sp. PCC 6803 (Schmidt *et al*., 2024). However, the extent to which ^m4^C occurrence in promoter sequences affects bacterial gene expression remains to be elucidated in a systematic way.

Cyanobacteria are the only prokaryotes conducting oxygenic photosynthesis making them ecologically relevant as globally important CO_2_ sinks (Flombaum *et al*., 2013). Their unique physiology and redox chemistry have also generated interest for producing various chemicals based on photosynthesis, using CO_2_ and solar energy (Hagemann and Hess, 2018; Klähn *et al*., 2024). Among cyanobacteria, DNA methylation has been best studied in the model strain *Synechocystis* 6803 on global level (Hu *et al*., 2018), or using specific mutants in genes encoding DNMTs (Scharnagl et al., 1998; Hagemann et al., 2018). Six DNMTs have been identified in *Synechocystis* 6803, of which five (M.Ssp6803I–V) are active and contribute to DNA methylation, while one is likely a pseudogene (M.Ssp6803VI) (Hagemann *et al*., 2018). Analysis of the five associated methylation motifs in chromosomal DNA revealed that the most prevalent is GGCC, occurring 38,512 times in the chromosome, followed by GATC, CGATCG, GAAGGC and the bipartite motif GGAN_7_TTGG/CCAAN_7_TCC with 29,852, 10,236, 1,317, and 1,181 occurrences, respectively (Hagemann *et al*., 2018). Close homologs of the *Synechocystis* 6803 DNMTs I-III are encoded in many other cyanobacterial genomes pointing at their important function in the cyanobacterial radiation (Elhai, 2015).

This study is focused on the DNMT M.Ssp6803II targeting the GGCC motif, because this methylation event is by far the most frequent in *Synechocystis* 6803. Furthermore, its absence due to deletion of the M.Ssp6803II-encoding gene *sll0729* had a massive impact on growth and pigmentation, which could be reversed by ectopic expression of the gene (Hagemann *et al*., 2018), and by frequently occurring suppressor mutations (Gärtner *et al*., 2019). The rapidly increasing presence of suppressor mutants in cell suspensions of the mutant Δ*sll0729* made it initially impossible to analyze the impact of GGCC methylation on *Synechocystis* 6803. But we then found that the pigmentation phenotype was mainly caused by the unbalanced expression of protoporphyrinogen IX oxidase encoded by *hemJ*, due to the missing methylation of the first C within the GGCC motif in the core *hemJ* (*slr1790*) promoter (Schmidt *et al*., 2024). In that study, a strain MP*slr1790*/Δ*sll0729* (here called MPΔ*sll0729)* was generated, in which the GGCC motif in the *hemJ* promoter was substituted by GGTC, which resulted in wild-type like growth and pigmentation, while genome-wide GGCC methylation remained completely absent (Schmidt *et al*., 2024). Utilizing this strain, here we aimed to analyze the impact of ^m4^C methylation on gene expression globally. The MPΔ*sll0729* strain with complete inactivation of *sll0729,* and the control strain expressing *hemJ* from an identical single-nt substitution in the P*_hemJ_* promoter (here called MP), were cultivated under different conditions and the transcriptomes were analyzed using genome-wide microarrays. The newly obtained data were analyzed together with gene expression data from mutant Δ*sll0729* and the complementation strain 2014 Com, which were generated previously (Hagemann *et al*., 2018; Gärtner *et al*., 2019). The analyses revealed the reproducible dysregulation of more than 300 genes in the absence of M.Ssp6803II, which was verified by qPCR experiments of genes that bear GGCC sites close to the respective transcription start sites (TSSs). Some of these genes encode proteins of crucial physiological relevance, pointing at the biological relevance of ^m4^C methylation for bacterial gene expression.

## 2 Results and Discussion

### 2.1 Effect of Different Light Intensities of Methylation-Dependent Gene Expression

Here we used the following strains: (i) Δ*sll0729;* (ii) Δ*sll0729* suppressor mutant; (iii) MPΔ*sll0729* (GGCC◊GGTC mutated *hemJ* promoter, Δ*sll0729,spc^R^,kan^R^*); (iv) MP (GGCC◊GGTC mutated *hemJ* promoter in wild type background, *spc^R^*); (v) *Synechocystis* sp. PCC 6803 PCC-M (wild type, WT); (vi) a Δ*sll0729* complementation mutant, expressing *sll0729* under control of a zinc-inducible P*_ziaA_* promoter from plasmid pVZ322 backbone (2014 Com) (**Table S1**, for details, see (Gärtner *et al*., 2019); (Schmidt *et al*., 2024)).

Gene expression of *Synechocystis* 6803 strains carrying the GGCC to GGTC modified P*_hemJ_* promoter (strains MPΔ*sll0729* and MP) were first compared to wild type (WT) under standard low light (LL) or after 30 min at high light (HL) conditions. The results showed the rather similar upregulation of many known HL-inducible genes in the three strains (**Figure 1**), consistent with reported HL effects on gene expression in *Synechocystis* 6803 (e.g., Hihara et al., 2001; Alvarenga-Lucius et al., 2026). The 50 most highly HL-induced genes in WT cells were also most highly upregulated in the strains MPΔ*sll0729* and MP. Among them we detected many genes such as *sbtA*, *ndhF3*, *ccmM* for elevated inorganic carbon uptake and concentration (Hagemann *et al*., 2021), *hliC* and *hliA* for the HL-induced proteins known to prevent ROS production from chlorophylls (Konert *et al*., 2022), and *flv2* and *flv4* for the flavodiiron proteins involved in the dissipation of excess light energy (Nikkanen *et al*., 2025). Furthermore, many genes encoding ribosomal proteins were induced, consistent with the growth stimulation by more light energy (**Figure 1**). The down-regulated genes also widely overlapped in the three strains, including genes for phycobiliproteins (*cpcC2, cpcB, cpcA*), photosystem proteins (*psaL, psaF, psaA, psbO, psbT*), and electron transport factors (*petJ, fed5*). Only small quantitative differences were noticed in the degree of induction or suppression of these genes between the compared strains. These results clearly indicated that GGCC-specific DNA methylation had only a minor impact on the overall HL acclimation response of *Synechocystis* 6803.

**Figure 1.**
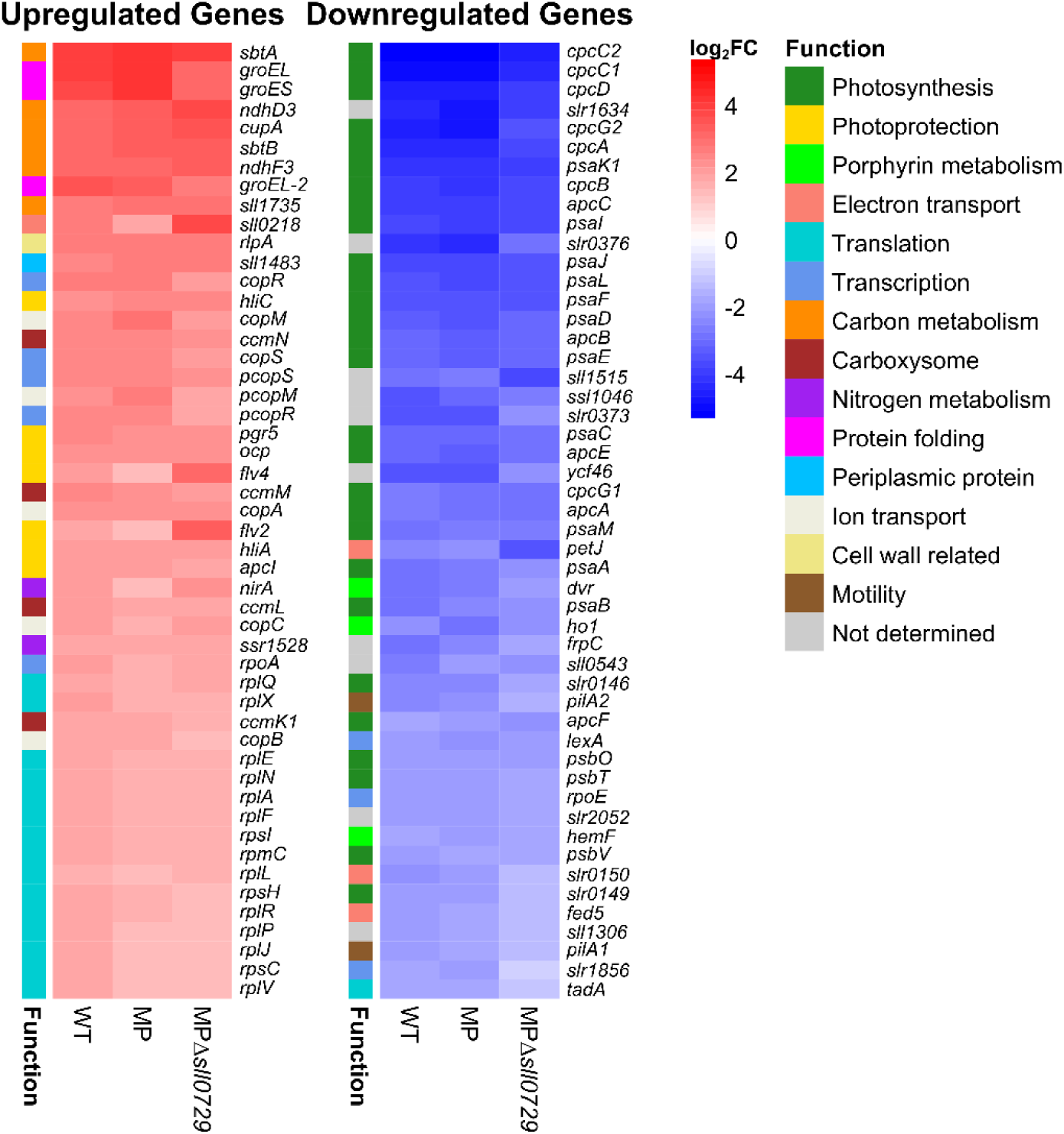
The 100 genes with most prominent changes in gene expression upon high light treatment. The wild type (WT), the control strain bearing the GGTC mutation in the *hemJ* promoter (MP), and the strain with the *hemJ* promoter mutation and deleted *sll0729* gene (MPΔ*sll0729*) were grown under standard low light (LL) and subjected to high light (HL) for 30 min. The left panel shows a heat map of the 50 most highly HL-induced genes, whereas the right panel shows the 50 most strongly down-regulated genes. Genes are grouped into different functional categories (Function).

However, despite the almost identical patterns of prominent gene expression changes after shifts from LL to HL conditions, we noticed a set of genes that were quantitatively differently regulated. Among them were the genes *slr1704*, *sll0470*, *sll0837*, *nblD* and *sigH,* which were more highly expressed in MPΔ*sll0729* than in the MP control strain. The opposite effect was observed for genes such as *argJ*, *sll1884*, *sll1305*, *slr1799* and *cupB* (**Figure 2**). These quantitative differences were observed under both, HL and LL conditions and indicate a GGCC-methylation-specific effect on their expression strength.

**Figure 2.**
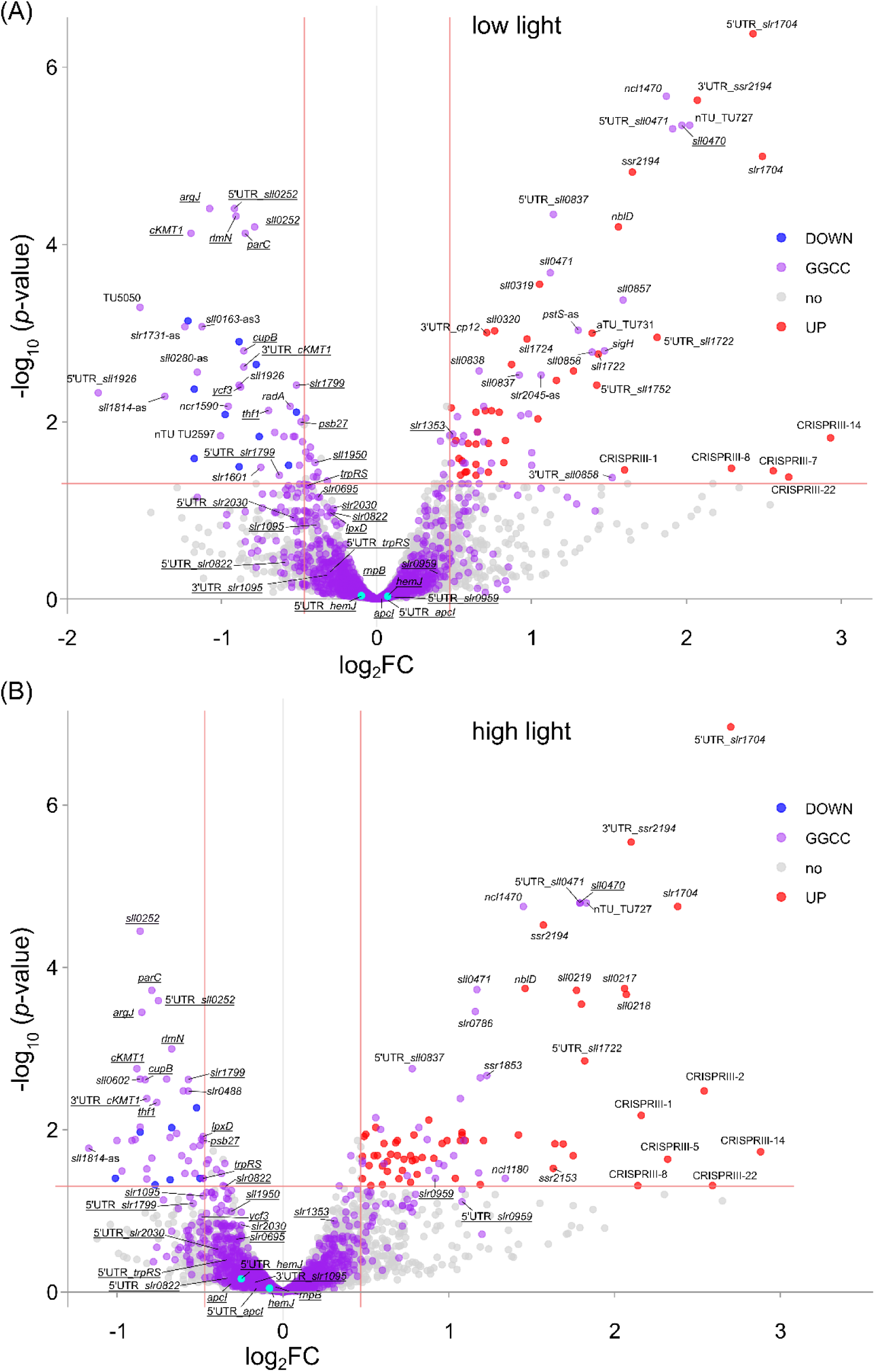
Transcriptomic differences depending on the presence (strain MP) or absence (strain MPΔ*sll072*9) of DNA GG^m^CC methylation. The volcano plots display log2FCs of differences in transcript levels between the two strains. **(A)** Comparison of transcriptome differences under standard low light (LL). **(B)** Differences after shift to high light (HL). Statistical significance is indicated by −log10 (*p*-value). Genes carrying a GGCC within their promoter sequence (−65 to +11) are displayed in purple, downregulated transcripts are colored in blue, upregulated transcripts in red, unaffected transcripts in light gray, qRT-PCR-validated transcripts are underlined. Threshold lines indicate cut-off values for significance. Transcripts are labeled by their gene names, gene IDs, or UTRs in case of annotated 5’ and 3’ UTRs. The CRISPR nomenclature follows Scholz *et al*. (2013).

Many of these genes contain a GGCC motif in their promoter region. Hence, this data set provides a solid basis for the specific selection of genes specifically influenced by ^m4^C methylation. To validate that these differences in transcript levels resulted indeed from the missing ^m4^C methylation, we took samples from the original Δ*sll0729* deletion mutant and the Δ*sll0729* suppressor mutant and analyzed selected genes by qRT-PCR (**Figure 3**). All genes except *sll0470* displayed lowered transcript levels in both the Δ*sll0729* mutant and its suppressor compared to WT. In the qPCR experiments, the most pronounced HL-specific changes in *Δsll0729* and Δ*sll0729*-suppressor compared to WT were observed for *parC*, *cKMT1*, *sll1950*, *apcI* and *gpx2*. It should be noted that increased transcript levels of *sll0470* and decreased transcript levels of *cKMT1* were observed previously and validated in promoter fusion experiments to be caused by altered transcription in the Δ*sll0729* mutant (Gärtner *et al*., 2019). Furthermore, the reported impact of the missing DNA methylation on *hemJ* transcript levels (Schmidt *et al*., 2024) was observed (**Figure 3**). Here, it was intriguing to see that the *hemJ* mRNA level in the Δ*sll0729* suppressor mutant was still lower relative to WT, although not as pronounced as in Δ*sll0729*. This points at the somewhat deviating regulation due to the substitution of cytosine by thymine in the discriminator sequence of the P*_hemJ_* promoter.

**Figure 3.**
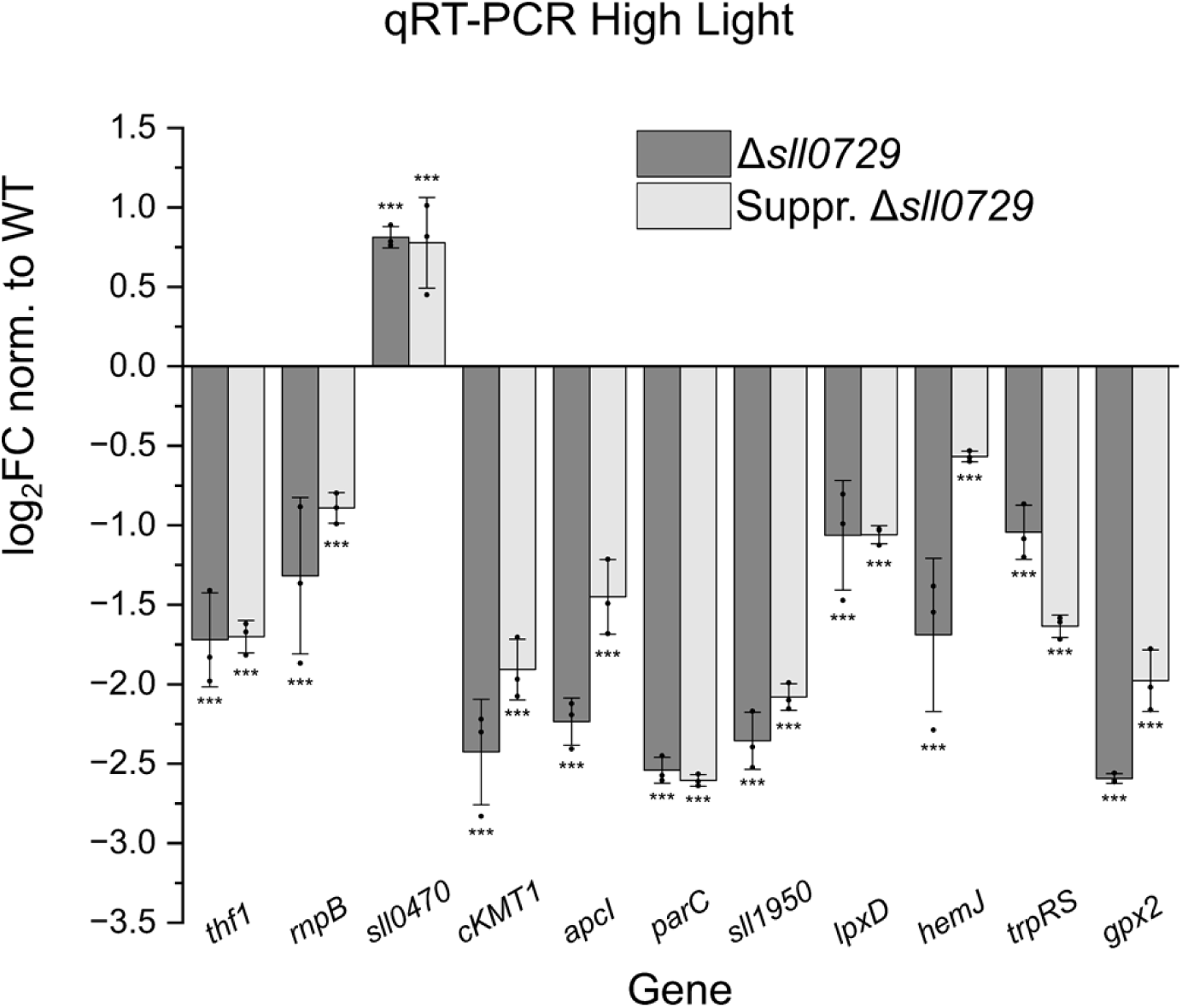
Gene expression levels in Δ*sll0729* (dark gray) and the Δ*sll0729* suppressor mutant (gray) relative to WT 30 min after shift to HL. Reverse transcription was performed using the Revertaid Kit. Expression levels are presented as log2FCs, normalized to WT. Bars represent the average of three biological replicates, each measured in technical triplicates (n = 9), with standard deviation (SD) shown. Significance was calculated with a two tailed t-test with unequal variance (Welch’s t-test; *P ≤ 0.05; **P < 0.01; ***P < 0.001). Cells were exposed to 200 µmol photons m^−2^ s^−1^ for 30 min.

#### 2.1.1 Data Mining Reveals Global Effects of Missing GGCC Methylation on Gene Expression

To assess the transcriptional impact of GGCC methylation globally, transcriptomic datasets comparing the gene expression of Δ*sll0729* mutants and WT were analyzed. These datasets were derived from two independent microarrays performed over a 10-year period (2014 and 2024). It is important to note that independently obtained Δ*sll0729* mutant strains were used to obtain transcriptomic data specific for the missing GGCC methylation avoiding interference due to accumulating suppressor mutant effects as discussed in Gärtner *et al*. (2019). In the transcriptomic dataset of 2014, a Δ*sll0729* mutant (2014 Ko) and a Δ*sll0729* complementation mutant, expressing *sll0729* under control of a zinc-inducible promoter (P*_ziaA_*) from the autonomous replicating plasmid pVZ322 (2014 Com), were compared under standard growth conditions. In the more recently performed microarray (2024), the strain bearing the GGCC◊GGTC mutation in the P*_hemJ_* promoter (MP) was compared to the mutant carrying the same point mutation in the Δ*sll0729* background (MPΔ*sll0729*) (Schmidt et al., 2024). The gene expression in these strains was compared under HL (2024 HL) and standard LL (2024 LL).

The rationale of this analyses was that genes with consistent log_2_FCs across all datasets except 2014 Com are likely to be directly regulated by GG^m4^CC methylation. Despite the variability of the data, a set of 408 transcripts showed differential expression across all datasets. Out of the 408 transcripts, 175 were consistently upregulated, whereas 233 were consistently downregulated in the Δ*sll0729* mutant and its derivatives compared to WT or strain MP. Genes involved in a wide range of cellular functions were found to be affected by the loss of GG^m4^CC methylation. To avoid loss of information, we intentionally didn’t set any *p*-value filter, as many genes did not meet conventional significance thresholds (≤0.05) due to variability between replicates. After this initial analysis, all transcriptomic datasets were reanalyzed requiring a |log_2_FC| ≥0.47 and *p*-value ≤0.05, and the expression differences across all datasets were visualized in heatmaps (**Figure 4**). While the magnitude of expression changes for most genes was modest, a consistent trend toward reduced transcript abundance was observed in the Δ*sll0729* mutants. This included genes encoding ribosomal proteins (*rps16*, *rpl36*, *rpl32*), the RNAP γ subunit (*rpoC1*), RNA-binding proteins (*rbp2* and *rbp3*), or *purC*. Notably, in the 2014 complementation strain (2014 Com), transcript levels of these genes were restored or showed a slight upregulation compared to WT (except *rbp2*). To further narrow down the number of potentially directly affected GGCC promoters, a threshold criterion ≥0.5 was applied. The datasets 2024 LL, 2024 HL, and 2014 Ko each were compared to 2014 Com (**Figure S1**). The comparison of consistently downregulated and upregulated genes lacking a GGCC motif in the core promoter is shown in **Figure S2.**

**Figure S1.**
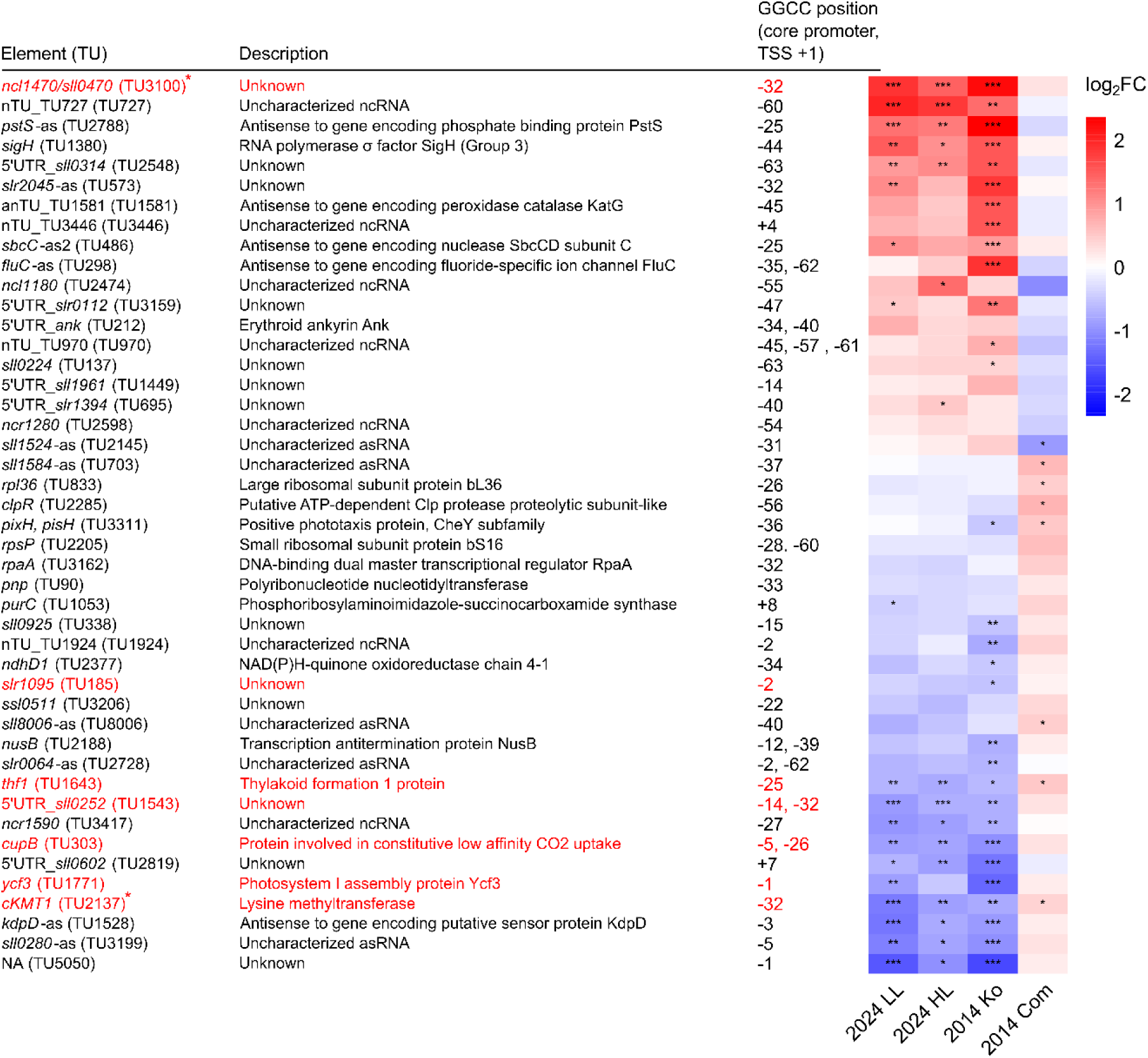
Heatmap of consistently downregulated and upregulated genes with GGCC in the core promoter sequence across multiple microarray datasets. The heatmap displays log2 fold changes in transcript levels between the WT or the MP control and different mutant strains from the years 2014 and 2024. In the left two columns, strains carrying the GGCC to GGTC modified P*hemJ* promoter in the Δ*sll0729* background (MP Δ*sll0729*) were compared to strain MP in the WT background, either cultivated at low light standard conditions (2024 LL), or after shift to high light for 30 min (2024 HL). In the right two columns, the Δ*sll0729* mutant (2014 Ko) or the Δ*sll0729* complementation strain (2014 Com) were compared to WT. Color scale ranges from blue (downregulated) to red (upregulated). Genes highlighted in red were selected for further analysis. ncRNA TU (nTU) ncRNA antisense to another TU not overlapping with the CDS (anTU). Numbers in brackets refer to TU number. Genes colored in red were validated via qRT-PCR and genes with an red asterisk were analyzed and confirmed by (Gärtner *et al*., 2019) Significance of the transcriptomic data was calculated with a two tailed t-test with unequal variance (Welch’s t-test; *P ≤0.05; **P <0.01; ***P <0.001). No significance criteria were used, but a log2 fold change criterion was applied, whereby each of the three microarray values (2024 LL, 2024 HL, and 2014 Ko) had to meet a difference ≥0.5 with 2014 Com. If more than one gene was part of a transcriptional unit, only the first gene is listed.

**Figure S2.**
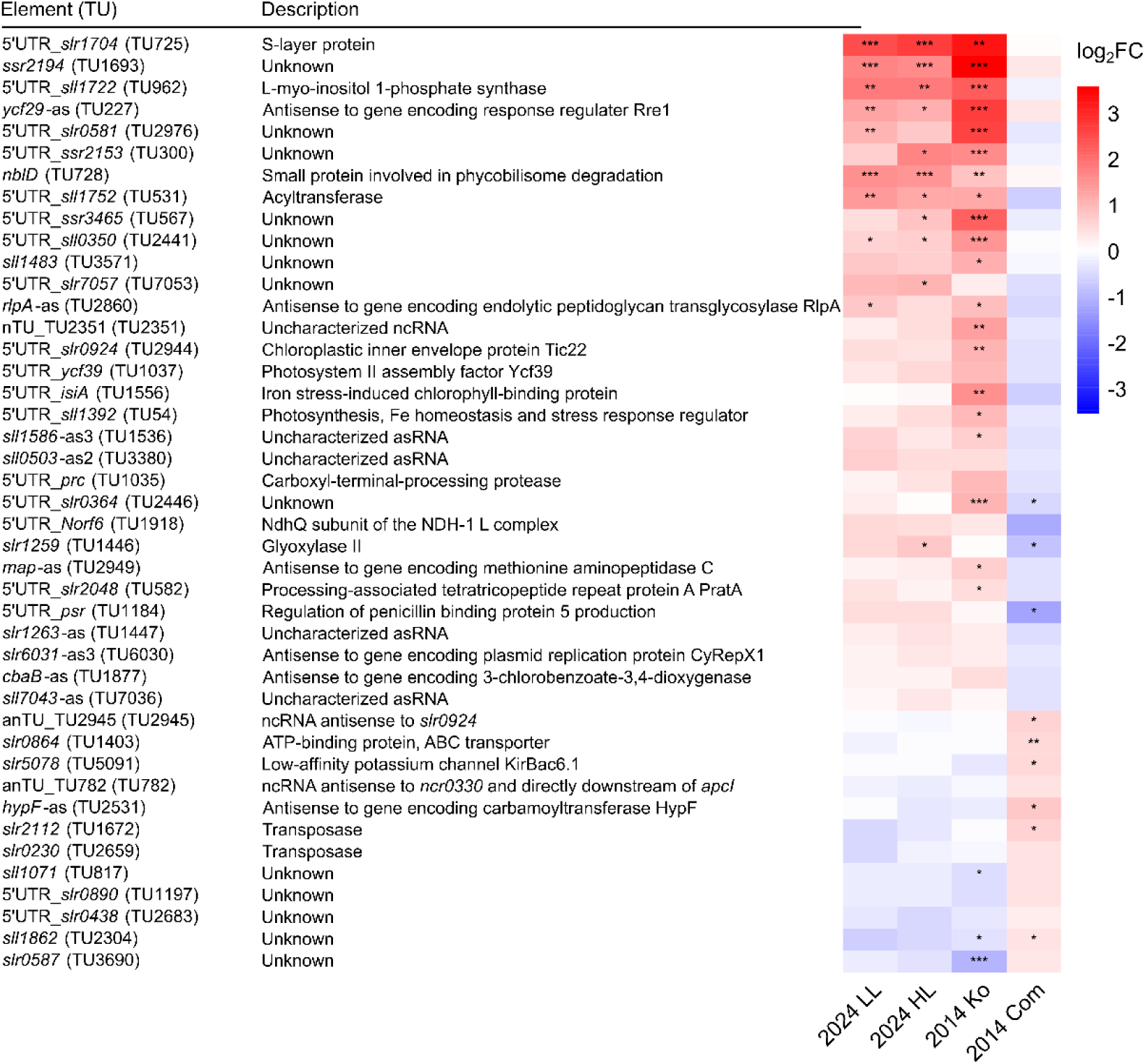
Heatmap of consistently downregulated and upregulated genes without a GGCC motif in the core promoter sequence across multiple microarray datasets. The heatmap displays log2 fold changes in transcript levels between the Δ*sll0729* mutants and WT strains from the years 2014 and 2024. Color scale ranges from blue (downregulated) to red (upregulated). Labels indicate the year of transcriptomic data acquisition, followed by the growth condition or strain: low light (LL) or high light (HL), Δ*sll0729* mutant (2014 Ko), and Δ*sll0729* complemented strain Δ*sll0729*::*0729* (2014 Com). Strains carrying the GGCC to GGTC modified P*hemJ* promoter in the Δ*sll0729* background (MP Δ*sll0729*) were compared to strain MP in the WT background, either cultivated at low light standard conditions (2024 LL), or after shift to high light for 30 min (2024 HL), a Δ*sll0729* mutant (2014 Ko), and a Δ*sll0729* complementation mutant (2014 Com), both compared to WT. ncRNA TU (nTU) ncRNA antisense to another TU not overlapping with the CDS (anTU). Numbers in brackets refer to TU number. Significance of the transcriptomic data was calculated with a two tailed t-test with unequal variance (Welch’s t-test; *P ≤0.05; **P <0.01; ***P <0.001). No significance criteria were used, but a log2 fold change criterion was applied, whereby each of the three microarray values (2024 LL, 2024 HL, and 2014 Ko) had to meet a difference ≥0.5 with 2014 Com. If more than one gene was part of a transcriptional unit, only the first gene was listed.

**Figure 4.**
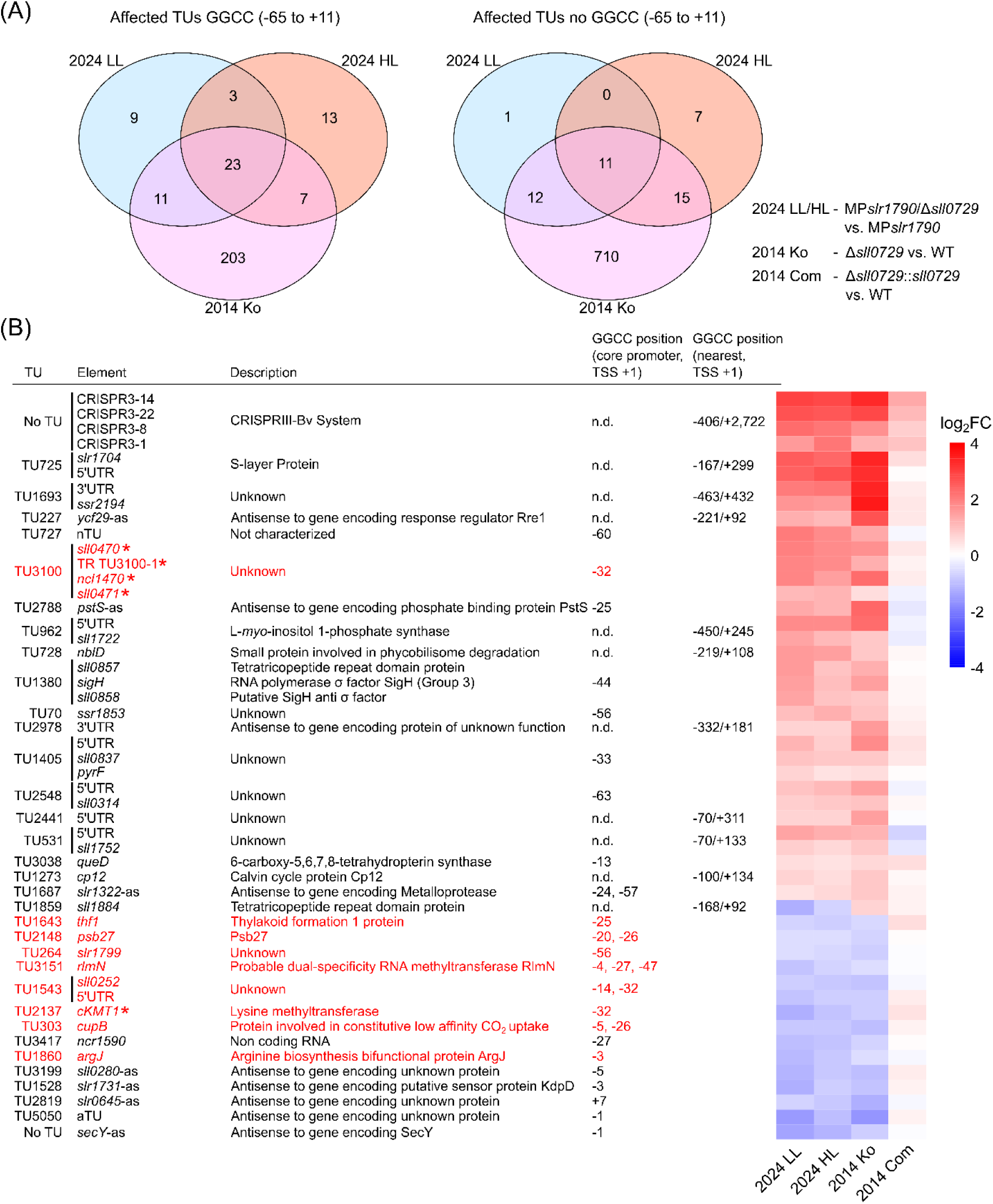
Significantly affected genes across three different transcriptomic datasets of Δ*sll0729* mutants. Strains carrying the GGCC to GGTC modified P*hemJ* promoter in the Δ*sll0729* background (MP Δ*sll0729*) were compared to strain MP in the WT background, either cultivated at low light standard conditions (2024 LL), or after shift to high light for 30 min (2024 HL), a Δ*sll0729* mutant (2014 Ko), and a Δ*sll0729* complementation mutant (2014 Com), both compared to WT. Transcriptomic datasets were obtained over a 10-year period. **(A)** Overlaps in the numbers of differentially expressed TUs, either carrying or lacking a GGCC motif within their promoter sequences spanning −65 to +11 (TSS+1). **(B)** Consistently up- or downregulated transcripts. The heatmap displays log2FC of transcript abundance comparing the same strains as in panel A. Only transcripts with a │log2FC│ ≤0.47 and *p*-values ≤0.05 are displayed. Transcripts are labeled by their TU numbers (Kopf *et al*., 2014), gene names, gene IDs, or UTRs in case of annotated 5’ and 3’ UTRs. The CRISPR nomenclature follows Scholz et al. (2013). Contiguous TUs are indicated by black vertical lines. Genes colored in red were validated via qRT-PCR and genes with an red asterisk were analyzed and confirmed by (Gärtner *et al*., 2019). The GGCC motif position is displayed regarding the position of the internal cytosine of the motif relative to the TSS (+1). (nTU) TU transcribing an ncRNA.

The overlap between all genes or transcripts showing either up- or downregulation in the Δ*sll0729* mutant strains relative to WT is visualized in **Figure 4A**. Across all datasets, a subset of 23 TUs carrying a GGCC motif within their promoter, as well as 11 TUs lacking this motif showed changes in transcript abundance, indicating a direct effect of ^m4^C loss rather than secondary effects. Nine of these genes (highlighted in red, **Error! Reference source not f ound.4B**), carrying a GGCC motif, were later validated by qRT-PCR (see below). Eight of the consistently deregulated transcripts (*sigH*, *slr1704,* TU727, *nblD*, *sll0314*, *sigH*, *sll0857,* and *sll0858*) were upregulated in the 2024 HL, LL, and 2014 Ko strains in Δ*sll0729,* while they showed unchanged transcript abundances in the complementation strain Δ*sll0729*::*0729* (2014 Com). TU1859, containing the gene *sll1884*, was downregulated in all Δ*sll0729* strains except for the 2014 Ko strain, where upregulated transcript levels were observed, while it was expressed in the Δ*sll0729* complementation strain 2014 Com like in WT (**Figure 4B**). TU1859 is lacking a GGCC motif within its promoter sequence. The closest GGCC motifs are located 92 bp downstream and 166 bp upstream regarding the TSS. Directly upstream of TU1859, TU1860 is positioned (*argJ*), which was one of the consistently affected genes across all five transcriptomic datasets and carries a GGCC in the discriminator sequence of its promoter (^m4^C position −3 relative to TSS). Therefore, the observed transcript abundance change could result from RNAP read-through from TU1860 into TU1859.

### 2.2 Validation of Gene Expression Changes by qPCR

For further analysis, a specific set of 26 genes carrying GGCC in functionally significant promoter elements were chosen and visualized as a separate heatmap (**Figure S3**). Those included genes encoding proteins e.g. the photosystem II lipoprotein Psb27 (*slr1645*) and probable dual-specificity RNA methyltransferase RlmN (*sll0098*), a ribozyme (*rnpB*), which is the RNA subunit of the ribonuclease P, and the regulatory small RNA (sRNA) *nsir4* involved in nitrogen metabolism (Klähn *et al*., 2015). These genes covered a variety of biological processes. As a control, *psbA2*, which lacks a GGCC motif in the defined −65 to +11 promoter window, was included. Additionally, the previously validated GGCC-methylation-sensitive promoters of *sll0470*, *cKMT1* (Gärtner *et al*., 2019), and *hemJ* (Schmidt et al., 2024) were used as internal controls for methylation-dependent regulation.

Most of the 26 genes showed consistently altered expression across all transcriptomic datasets. The genes *cupB*, *psb27*, *thf1*, *rlmN* (*sll0098*), *sll0252*, *cKMT1* (Gärtner *et al*., 2019), *argJ*, *parC* (*sll1941*), *lpxD* (*slr0776*), *slr0822*, *ycf3* (*slr0823*), *slr1095*, *hemJ* (*slr1790*), *trpRS* (*slr1884*), *gpx2* (*slr1992*), *nsir4*, *slr1799* and *slr2030* were consistently downregulated across experimental datasets. The most pronounced decreases in transcript abundance were detected for *apcI* and *gpx2* in the 2014 datasets (log_2_FCs −2.73 and −1.88). Among upregulated genes, *sll0470* exhibited the highest expression changes, with log_2_FC values of 2.25 in the 2014 Δ*sll0729* mutant (2014 Ko) and 1.87 in the MPΔ*sll0729* mutant under HL conditions (2024 HL). All upregulated genes *sll0470*, *slr1353*, and *slr0959* displayed consistent log_2_FCs and either inverse or unchanged (but reduced to WT similar levels) patterns in the Δ*sll0729* complementation strain (2014 Com) compared to WT and other strains.

Several genes did not show consistently altered expression across all transcriptomic datasets. Specifically, the genes *rnpB*, *apcI* (*sll1911*), *sll1950* and *slr0695* revealed minor fluctuations in gene expression in one (*rnpB*, *apcI*, *sll1950*) dataset, contrary to their overall expression trends. The gene *apcI*, encoding a phycobilisome linker protein, was unaffected under LL conditions (2024 LL) but downregulated in all other strains. However, native transcript levels were not restored to WT conditions in the *sll0729* complementation mutant (2014 Com) suggesting a ^m4^C independent regulation of *apcI* transcription. Furthermore, *psbA2*, *rnpB* and *hemJ* showed no differential expression in 2024 for HL and LL conditions when comparing MP to MPΔ*sll0729*, but in the other transcriptomic datasets. The absence of differential expression for *hemJ* confirmed the restored transcription following the substitution of cytosine by thymine at the -4 position (TSS +1). Hence, for *rnpB* and *psbA2*, altered transcription may occur indirectly as a downstream effect of *hemJ* regulation in transcriptomic datasets 2014 Ko. Notably, *psbA2* expression was different in Δ*sll0729* mutants despite the lack of a GGCC recognition motif in its promoter. The nearest GGCC motifs are located 270 nt upstream and 881 nt downstream of the TSS of *psbA2*. Therefore, these genes did not meet the requirements to be displayed in the heatmap (**Figure 4** and **Figure S1**), which visualized only those genes exhibiting uniform expression changes across the 10-year period.

**Figure S3.**
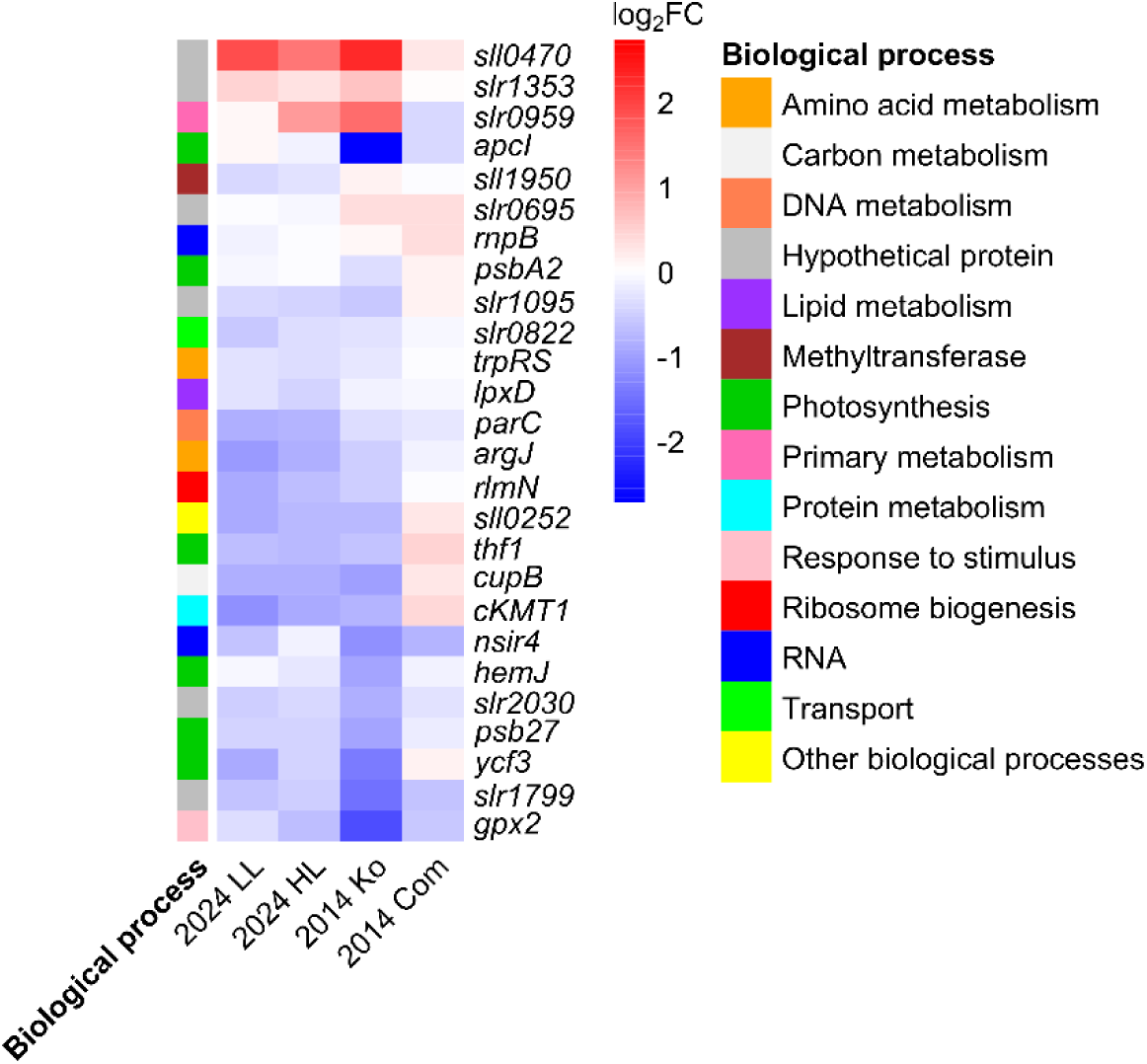
Heatmap of 26 genes from transcriptomic datasets collected over a 10-year period. Each gene’s promoter region contains a GGCC motif within the −65 to +11 nt window relative to the TSS (+1). The heatmap displays log2FCs in gene expression for the same strains as in Figure 4B. The color gradient ranges from blue (downregulated) to red (upregulated). Colored squares next to the heatmap indicate the associated biological process with the specific gene.

To validate transcriptomic findings, qRT-PCR was performed. Here, the Δ*sll0729* mutant was freshly generated, together with the other strains. Therefore, validated transcript levels support the assumption that loss of ^m4^C in these promoter sequences causes a consistent effect in the transcription of these genes. Overall, qRT-PCR results were in agreement with microarray data for all genes (**Figure 5**). From these data we conclude that the consistency of the microarray-derived transcriptome (**Figure S1** and **S2**) and the qRT-PCR data (**Figure 5**) show that the changes in gene expression were caused by the lack of methylation and not a consequence of *hemJ*-dysregulation.

**Figure 5.**
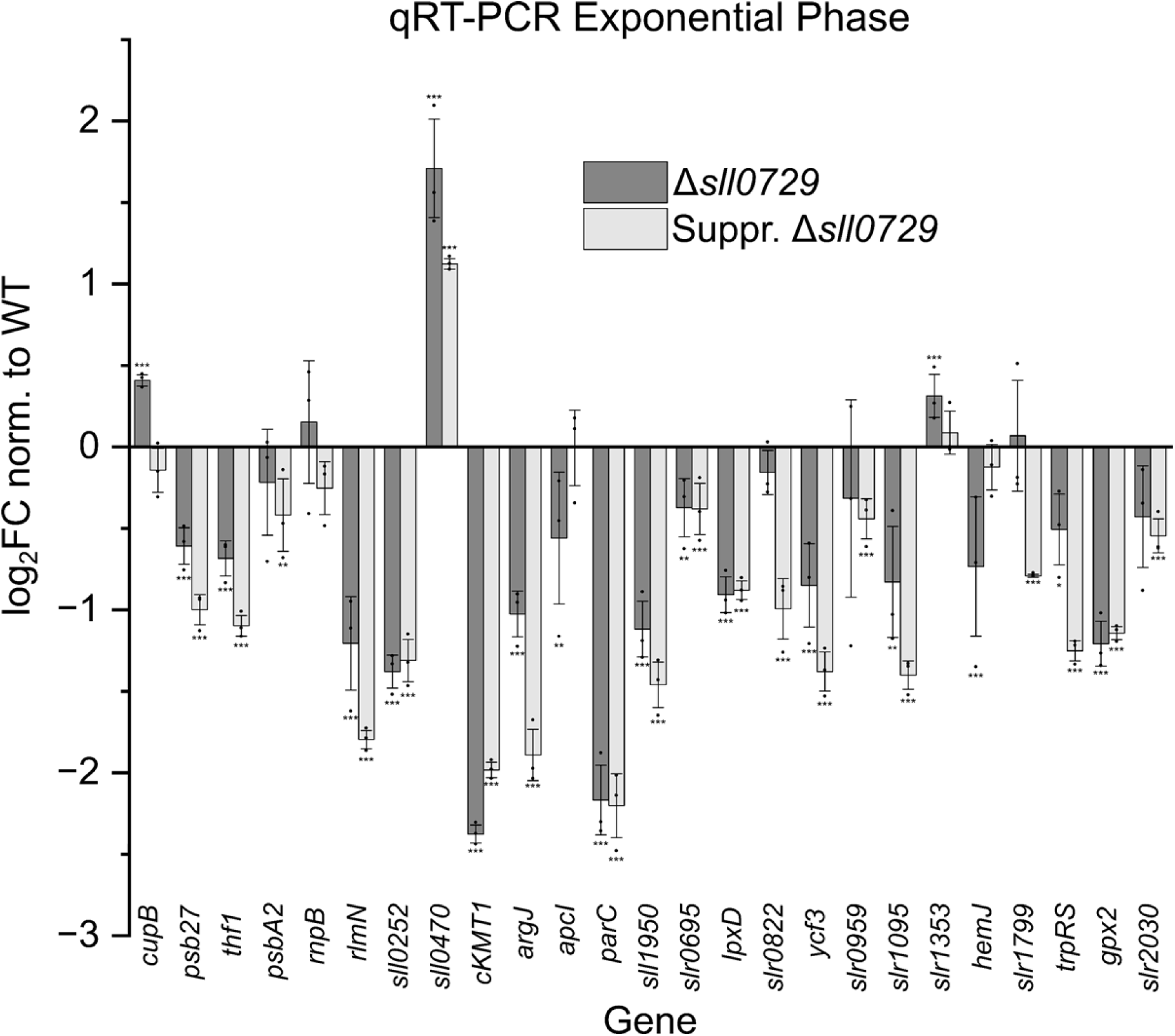
Gene expression level in the Δ*sll0729* mutant (dark gray) and the suppressor mutant Δ*sll0729* (gray) relative to WT during exponential phase. Reverse transcription was performed using the RevertAid Kit. Expression levels are presented as log2FCs, normalized to WT. Bars represent the average of three biological replicates, each measured in technical triplicates (n = 9), with standard deviation shown. Significance was calculated with a two tailed t-test with unequal variance (Welch’s t-test; *P ≤0.05; **P < 0.01; ***P < 0.001).

Furthermore, Northern hybridizations performed for *cupB*, *psb27*, and *thf1* (**Figure S4**) were consistent with both the transcriptomic and the qRT-PCR data. These results reinforce the reliability of the observed gene expression alterations associated with the *sll0729* deletion.

**Figure S4.**
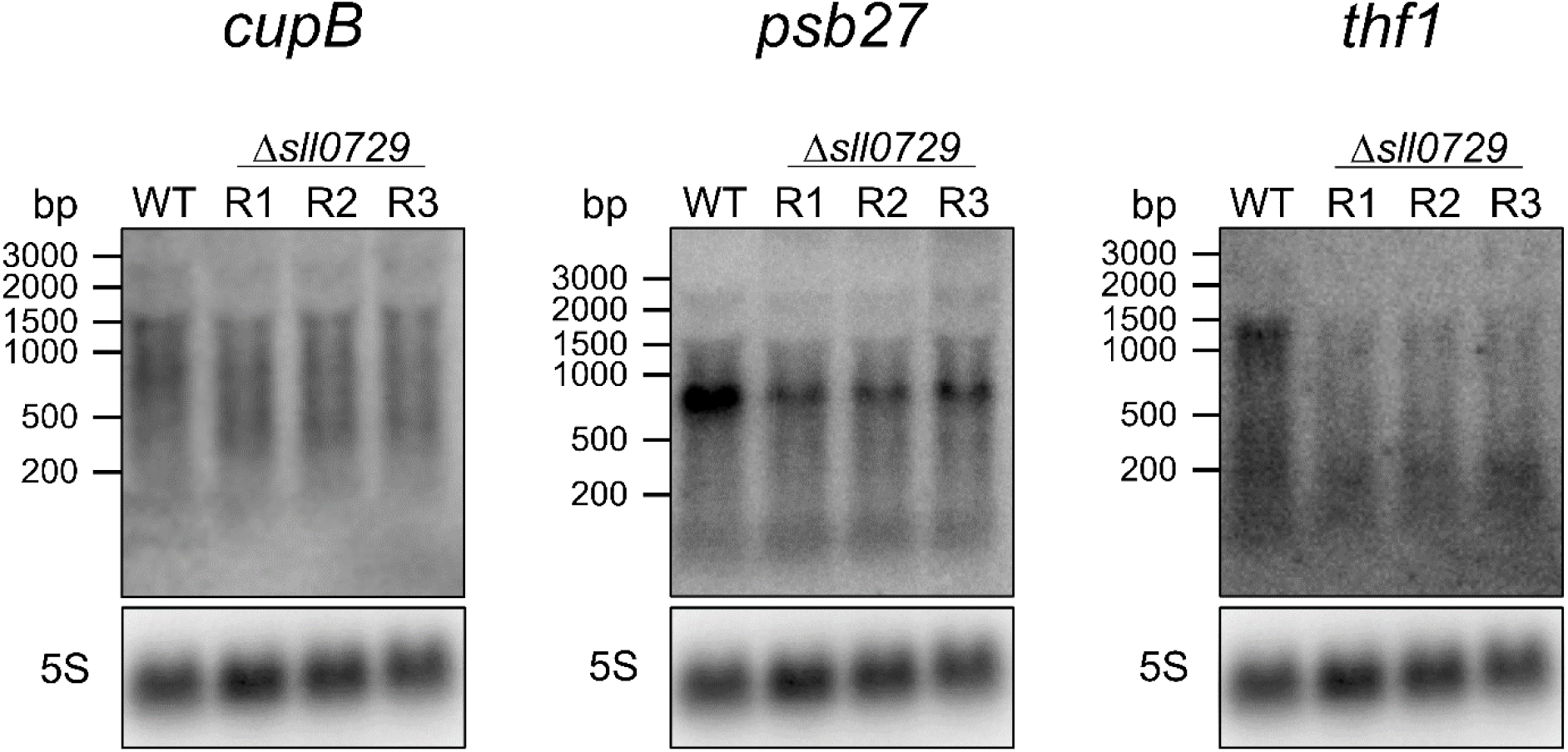
Northern hybridizations of *cupB*, *psb27* and *thf1* transcripts. Transcript levels in WT and three biological replicates of the Δ*sll0729* mutant (R1, R2 and R3) were compared. RNA levels were normalized to 5S rRNA as a loading control. Transcript sizes are indicated in base pairs (bp). Blots represent replicate experiments using equally treated RNA samples extracted at different time points.

### 2.3 Classification of Consistently Dysregulated Promoter Sequences

To get a deeper understanding of the methylation-sensitive promoters, a promoter classification was performed. Promoters were categorized into three different types as defined by Asayama and Imamura (2008) for *Synechocystis* 6803 according to their respective σ factor: **(i)** Promoters recognized by the group 1 “housekeeping” σ factor SigA (σ^A^, *slr0653*), **(ii)** the group 2 stress response σ factors SigB (σ^B^, *sll0306*), SigC (σ^C^, Sll0184), SigD (σ^D^, Sll2012) and SigE (σ^E^, Sll1689), **(iii)** the group 3 σ factors SigF (σ^F^, Slr1564), SigH (σ^H^, Sll0856). We did not classify group 4 promoters dependent on SigG (σ^G^, Slr1545) or SigI (σ^I^, Sll0687) (Srivastava *et al*., 2020), because these are not well characterized.

Characteristics of the different promoter sequences were used for the *in-silico* classification of the 26 differentially regulated genes (**Figure S3**). Promoter sequences that carried either TTG as the first three nt, or at least 4 matching nt of the −35 element consensus sequence TTGACA within the range of −28 to −37 were extracted as potential Type 1 promoters. Moreover, they had to fulfill the criteria of a spacer range (between the −10 and −35 element) of 16 to 18 nt. All other promoters were defined as Type 2 promoters. Furthermore, Type 2 promoters were screened for the presence of the canonical NtcA binding site (GTA–N_8_–TAC) or the degenerated motif GY–N_9-10_–AC, as a representative example of transcription factor binding sites. This analysis aimed to assess whether overlap with a GGCC motif was associated with altered transcript levels when comparing standard growth conditions with nitrogen starvation in the Δ*sll0729* mutant relative to WT.

Regarding the classification of Type 3 promoters by (Imamura and Asayama, 2009), the spacer distance between the −12 and −32 element spans approximately 15 nt. Moreover, the distance between the −12 element and the TSS can vary from 6 to 18 nt (Imamura and Asayama, 2009). Therefore, all promoter sequences were searched for characteristic Type 3 promoter elements, regardless of their distance from the TSS. First, a sequence similar to the −32 element was searched matching at least 3 nt of the characteristic −32 sequence TAGGC. Secondly, a −12 element matching the sequence GGTA was searched allowing a distance of 14 to 16 nt downstream of the −32 element.

Promoter sequences of eight genes were classified as Type 1 promoters and the remaining as Type 2 promoters. The promoter sequences of genes *psb27*, *slr2030* and *slr0695* did reveal potential Type 3 promoter elements (**Figure 6**). This search was also done for the 232 consistently regulated genes in the Δ*sll0729* mutants. Overall, 18.9% of the promoter sequences were classified as Type 1 and 81.1% as Type 2. Approximately 7% of the predicted Type 1 and Type 2 promoters did harbor sequence elements qualifying them as possible Type 3 promoters.

**Figure 6.**
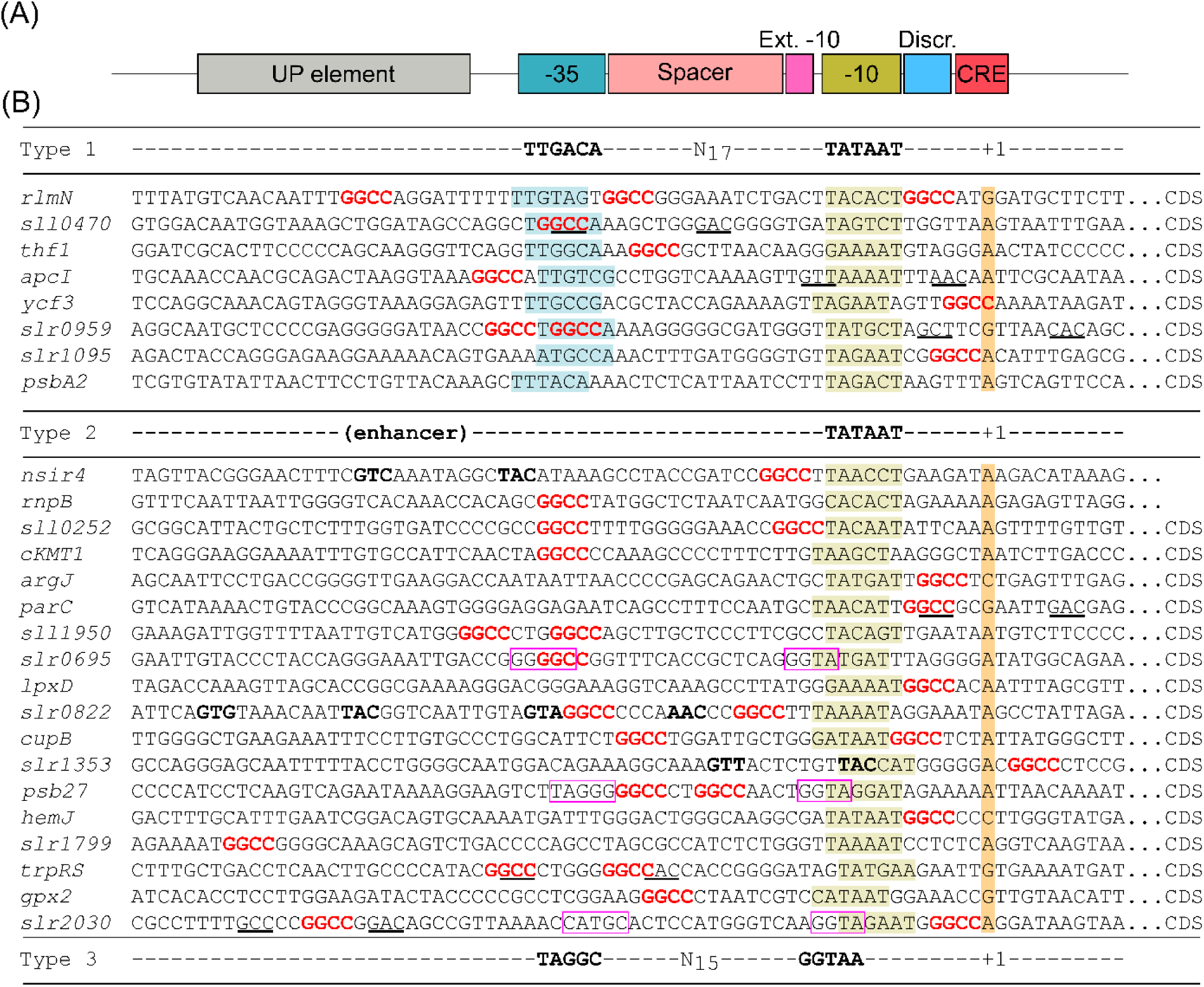
Nucleotide alignment of 26 promoter sequences containing a GGCC motif. **(A)** Schematic representation of promoter elements corresponding to the aligned promoter sequences shown in (B). **(B)** Promoter regions span from −65 to +11 relative to the TSS (+1). The −10 element is marked with a dark yellow box, the −35 element with a turquoise box, and the TSS is highlighted with an orange box. Already predicted −10 elements were used and new −10 elements were predicted using the PSWM of Mitschke et al. (2011). GGCC motifs are shown in bold red. Predicted NtcA-binding sites with high similarity to the canonical GTA–N8–TAC motif (Giner-Lamia *et al*., 2017) are highlighted in bold letters, while those with lower similarity are underlined. Type 3 promoter elements are indicated with a purple rectangle. Discr., discriminator region; CRE, core recognition element; CDS, coding sequences.

The distribution of GGCC motifs within extracted promoter sequences was examined. Histogram analysis revealed an almost complete absence of GGCC motifs within the segment containing the −10 element, while it was otherwise equally distributed. An exception was noticed for +1 (TSS) and +3 relative to the TSS. Here, an almost complete exclusion of the GGCC motif was observed (**Figure S5**).

**Figure S5.**
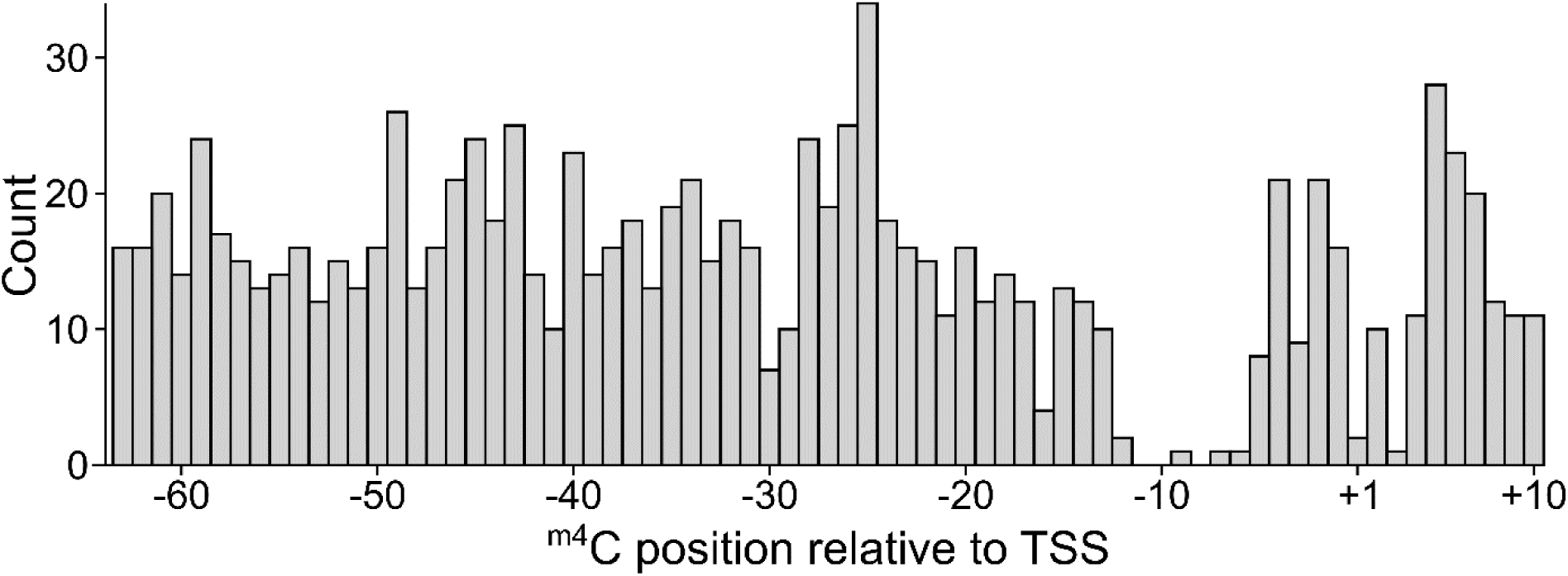
Histogram of GGCC motif distributions within chromosomal promoter sequences relative to the TSS (+1). Promoter sequences were defined as the −65 to +11 nt relative to the TSS. The histogram displays the position of the internal cytosine within each GGCC motif, resulting in a slightly shorter effective distance than the full promoter range.

To investigate positional effects of ^m4^C within GG^m4^CC motifs on transcription, data from three independent transcriptomic microarray experiments were reanalyzed. Annotated TUs (Kopf *et al*., 2014) containing at least one GGCC motif within the defined promoter window (−65 to +11 relative to TSS +1) were selected. For each transcriptomic dataset, log_2_FC values were plotted against the position of the internal C of the GGCC motif within promoters using scatter plots, with trends visualized via locally weighted scatter plot smoothing (loess regression (Cleveland and Devlin, 1988)). Expression values were normalized to WT as a calibrator strain. The 2024 LL and HL datasets, along with the 2014 complementation strain (2014 Com), were directly compared (**Figure 7**). Analysis for the 2014 Ko dataset was also performed and showed the same results (**Error! Reference source not found.**). Promoter sequences carried between o ne and three GGCC motifs (**Figure 7**). A consistent and clear downregulation was observed for promoters carrying a GGCC motif within the discriminator sequence region (**Figure 7A, B**).

**Figure 7.**
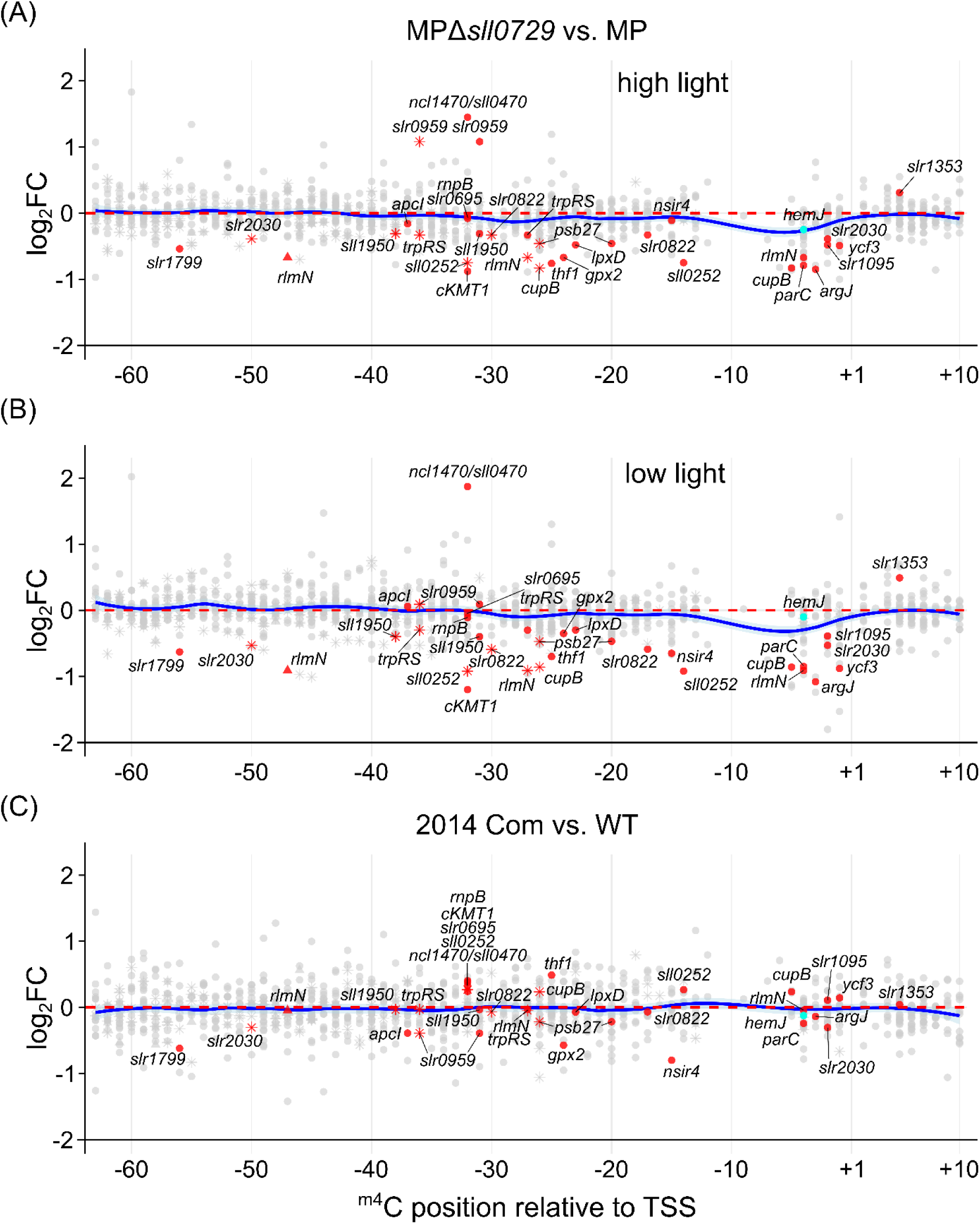
Expression of genes carrying a GGCC in their promoter sequences relative to the TSS (+1) of the Δ*sll0729* mutant. Each data point represents the log2FC in gene expression in Δ*sll0729* vs WT plotted against the position of the internal cytosine of the GGCC motif within the defined promoter window (−65 to +11 relative to the TSS). qRT-PCR analyzed transcripts are colored in red and *hemJ* in cyan. **(A)** *Synechocystis* strains carrying the GGCC to GGTC modified P*slr1790* promoter (MP) in the methyltransferase *sll0729* knockout background (MPΔ*sll0729*) compared to MP in the WT background after shift to LL for 30 min. **(B)** After shift to HL for 30 min. **(C)** *Synechocystis* Δ*sll0729* complemented knockout mutant (2014 Com) compared to WT background. A less smoothed trend line was calculated (blue line) showing the trend of the expression change depending on the location of the GGCC. Motif occurrences were denoted relative to the TSS as follows: first occurrence (●), second occurrence (**✳**), third occurrence (▴).

In contrast, promoters with GGCC motifs in other key sequence elements exhibited more variable transcriptional patterns. Promoters carrying a GGCC in the −35 element were both up- and downregulated across datasets. Motifs occurring downstream of the −35 element were associated with a mild decrease in transcript abundance (log_2_FC slightly below 0), followed by a decrease to log_2_FC of −0.5 reaching the discriminator sequence and a recovery toward baseline expression levels at the TSS (+1). For the UP-element region, no distinct trend in transcriptional regulation associated with GGCC positions was detected. Most promoter sequences harboring a GGCC motif appeared unaffected by the loss of ^m4^C methylation. If methylation was restored in the Δ*sll0729* complementation strain 2014 Com, the tendency to downregulated promoters carrying a GGCC motif within the discriminator sequence was not observed anymore, confirming the relevance of this modification in this position (**Figure 7C**). For promoters with the GGCC motif in other positions, different effects were observed in the Δ*sll0729* complementation strain: some gene promoters, e.g. *sll0252* and *cKMT1* showed an inverse trend with positive log_2_FCs, while *slr1799*, *gpx2*, and *slr2030*, remained unaffected or showed only minimal transcript variation upon complementation.

## 3 Conclusion

We previously demonstrated the functional relevance of GG^m4^CC methylation in the discriminator region for the proper expression of *hemJ* and the precise control of tetrapyrrole biosynthesis in *Synechocystis* 6803 (Schmidt *et al*., 2024). This raised the question if GG^m4^CC methylation could be relevant also for the expression of other genes. While previous analysis focused solely on the impact of ^m4^C on transcription initiation, we here considered all genes by examining transcript abundances at the TSS/5′UTR of each TU. Affected genes were sorted whether a GGCC was located within the promoter sequence spanning region −65 to +11 regarding the TSS, or not. 23 and 11 TUs were found to be significantly affected in all datasets carrying a GGCC and not, respectively. Overall, the 2014 strains showed a higher number of affected genes compared to the 2024 strains. Further, the strains of 2014 showed a higher number of affected genes lacking a GGCC motif than genes carrying a GGCC motif within their promoter sequence. However, it should be noticed that the Δ*sll0729* mutant culture in 2014 certainly contained already some suppressor mutant cells making this data set less reliable compared to 2024. Whereas for the 2024 strains almost similar numbers of genes were affected for both categories. We conclude that, compared to the relevance for *hemJ* expression, GG^m4^CC DNA methylation impacts the transcription of far more genes than previously anticipated, although mostly at subcritical levels. Nevertheless, we identified several physiologically highly relevant genes involved in photosynthesis or the uptake of inorganic carbon to belong to the affected genes, raising an exciting topic for further research.

**Figure S6.**
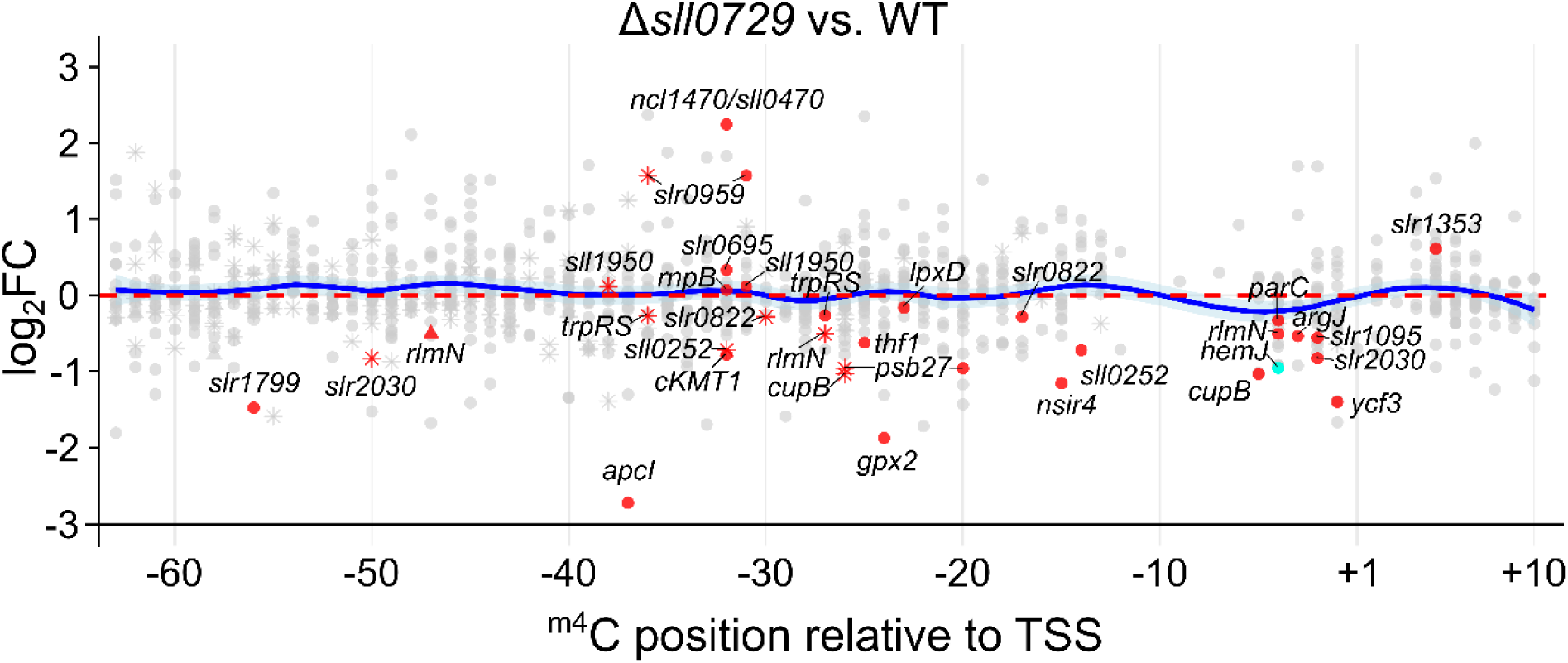
Expression of genes carrying a GGCC in their promoter sequences relative to the TSS (+1) of the Δ*sll0729* mutant. Each data point represents the log2FC in gene expression in Δ*sll0729* vs WT plotted against the position of the internal cytosine of the GGCC motif within the defined promoter window (−65 to +11 relative to the TSS). The transcriptomic data of 2014 is displayed. qRT-PCR analyzed transcripts are colored in red and *hemJ* in cyan. A loess smoothing was calculated (blue line) showing the trend of the expression change depending on the location of the GGCC. Motif occurrences were denoted relative to the TSS as follows: first occurrence (●), second occurrence (**✳**), third occurrence (▴).

## 4 Experimental Procedures

### 4.1 Strains and Growth Conditions

*Synechocystis* sp. PCC 6803 PCC-M (Trautmann *et al*., 2012) served as wild type (WT). Axenic strains were maintained on agar plates with BG11 mineral medium at 30°C under constant illumination (30 µmol photons m^-2^ s^-1^; to slow down formation of suppressor clones). Mutant strains of the methyltransferase M.Ssp6803II of *Synechocystis* 6803 (Δ*sll0729*) as well as complementation strains complementing the deletion of *sll0729* (Δ*sll0729::sll0729* complementation mutant, 2014 Com) were already described by (Hagemann *et al*., 2018; Gärtner *et al*., 2019; Schmidt *et al*., 2024). All strains are listed in **Table S1**.

For physiological characterization, axenic cultures of the different strains were grown photoautotrophically in BG11 medium (Rippka *et al*., 1979), either with gentle shaking in Erlenmeyer flasks at 30 µmol photons m^-2^ s^-1^, or aerated with ambient air in a photobioreactor (Multi-Cultivator MC 1000-OD-WW warm light, Photon Systems Instruments) at 30°C under continuous illumination of 40 µmol photons m^-2^ s^-1^ (standard conditions). For strain maintenance, antibiotics were included depending on the strain that was used (Schmidt *et al*., 2024). For experiments, the cultures were washed three or four times to deplete antibiotics in the medium and resuspended in fresh BG-11 medium without antibiotics. The time points of harvesting cell cultures were chosen according to the needed OD. For HL conditions, cells were exposed to 200 µmol photons m^−2^ s^−1^ once they reached an OD_750_ of 0.8. After 30 min of HL exposure, cells were collected for RNA extraction by filtration.

### 4.2 RNA Extraction and qRT-PCR

Cells from liquid cultures (OD_750_ ∼0.8) were harvested by vacuum filtration on hydrophilic polyethersulfone filters (Pall Supor®-800, 0.8 µm). Filters were immersed in 1 mL PGTX (Pinto *et al*., 2009), snap frozen in liquid nitrogen and stored at −80 °C until further processing. Frozen cultures were incubated for 15 min at 65 °C. 700 µl of chloroform/isoamyl alcohol (IAA) (24:1) was added, mixed and incubated for 10 min at room temperature (RT). During incubation samples were vortexed several times. Phase separation was performed by centrifugation (3,260 × g, 20 °C, 3 min). The upper aqueous phase was transferred to a new tube and mixed with one volume of chloroform/IAA (24:1). Phase separation and the chloroform/IAA washing step were repeated. After the second chloroform/IAA washing step, the aqueous upper phase was transferred to a new tube, mixed with an equal volume of isopropanol and precipitated overnight (ON) at −20 °C. The precipitated RNA was pelleted by centrifugation (12,000 × g, 4°C, 30 min). The isopropanol was discarded, and the pelleted RNA was washed with 200 µl of 70% ethanol. The RNA was centrifuged (12,000 × g, 4 °C, 10 min) and the ethanol was discarded. The RNA pellet was air-dried and resuspended in 50 to 100 µl of nuclease-free water. RNA concentration was determined using a spectrophotometer (NanoDrop ND-1000, Peqlab) and a fluorometer (Invitrogen™ Qubit™ 3). Quality control and RNA concentration were assessed with denaturing gel electrophoresis.

Quantitative reverse transcription polymerase chain reaction (qRT-PCR) was performed using the FastGene qFYR Real-Time PCR System (NIPPON Genetics). cDNA synthesis was performed using the RevertAid First Strand cDNA Synthesis Kit (Thermo Scientific™) according to the manufacturer’s instructions using 600 ng of each RNA sample as template. Prior cDNA synthesis RNA was DNase treated to reduce background amplification. DNase treatment was performed using the TURBO DNA-free™ Kit (Invitrogen™) according to the manufacturer’s protocol. Random hexamer and target specific primers (**Table S2**) were used for cDNA synthesis. Primer matrices were tested to obtain useful concentrations for the qPCR amplification. PCR amplification programs followed these steps: Hold Stage (3 min, 95 °C), PCR Stage ([5 s, 95 °C followed by 20 s, 62 °C] × 40), and Melt Curve Stage (15 s, 95 °C, 1 min, 60 °C -> 95 °C in 0.2 °C steps). The qPCR mix qPCRBIO SyGreen® Blue Mix separate-ROX (PCR Biosystems) was used PCR reactions were performed according to PCR Biosystems protocol. Primers were synthesized by Integrated DNA Technologies, Inc. (IDT). All reactions were measured as technical triplicates and RNase P subunit A (*rnpA*) was used as an internal control. The data were analyzed using the qFYR Analyzer Studio Software (Nippon Genetics) and relative quantification of RNA was calculated using the comparative Ct Method (ΔΔCt) using WT sample as the calibrator.

### 4.3 Expression Analyses

For microarray analyses, RNA samples from two to three biological replicates were hybridized to Agilent 8×44K microarrays following published sample preparation and hybridization procedures (Voß and Hess, 2014). Briefly, 2 µg of DNase-treated RNA was labeled with Cy3 using ULS Fluorescent labeling Kit for Agilent Arrays (Kreatech). Microarray hybridization was performed with 600 ng of Cy3-labeled RNA for 17 h at 65 °C. Microarray raw data were acquired using an Agilent Microarray Scanner C (model G2505C) and processed with Agilent Scan Control and Feature Extraction software (version 10.7.3.1). Data analysis was performed using the limma R package (version 3.52.4) (Ritchie *et al*., 2015). Data were normalized using the R-implementation in limma (Ritchie *et al*., 2015), and p values derived from t-tests were adjusted for multiple testing using Benjamini-Hochberg method to control the false discovery rate (Benjamini and Hochberg, 1995).

For Northern Blots, 5 µg of RNA were separated by size on 1.5% formaldehyde-agarose gels and blotted onto a Hybond-N+ membranes (Amersham) by capillary transfer. Northern hybridizations were performed using single-stranded isotope-labeled RNA probes transcribed from PCR-generated templates *in vitro* (for oligonucleotide sequences, see **Table S2**). Radioactively labeled probes were generated using [α-^32^P]-UTP (Hartmann Analytic) and the Maxiscript™ T7 *In vitro* transcription kit (Thermo Fisher Scientific). Hybridizations were performed in hybridization buffer (0.25 M NaCl, 0.12 M NaP_i_ buffer p*H* 7.2, 7 % SDS (v/v)) at 62 °C. After washing, membranes were re-hybridized with an end-labeled 5S rRNA oligonucleotide probe. Polynucleotide kinase (Thermo Fisher Scientific) was used for end-labeling with [γ-^32^P]-ATP (Hartmann Analytic). Signals were detected with a Laser Scanner Typhoon FLA 9500 (GE Healthcare) using Quantity One software and normalized to the signal of the 5S rRNA.

### 4.4 Bioinformatic Analyses

The bioinformatic workflow is depicted in **Figure 8**.

**Figure 8.**
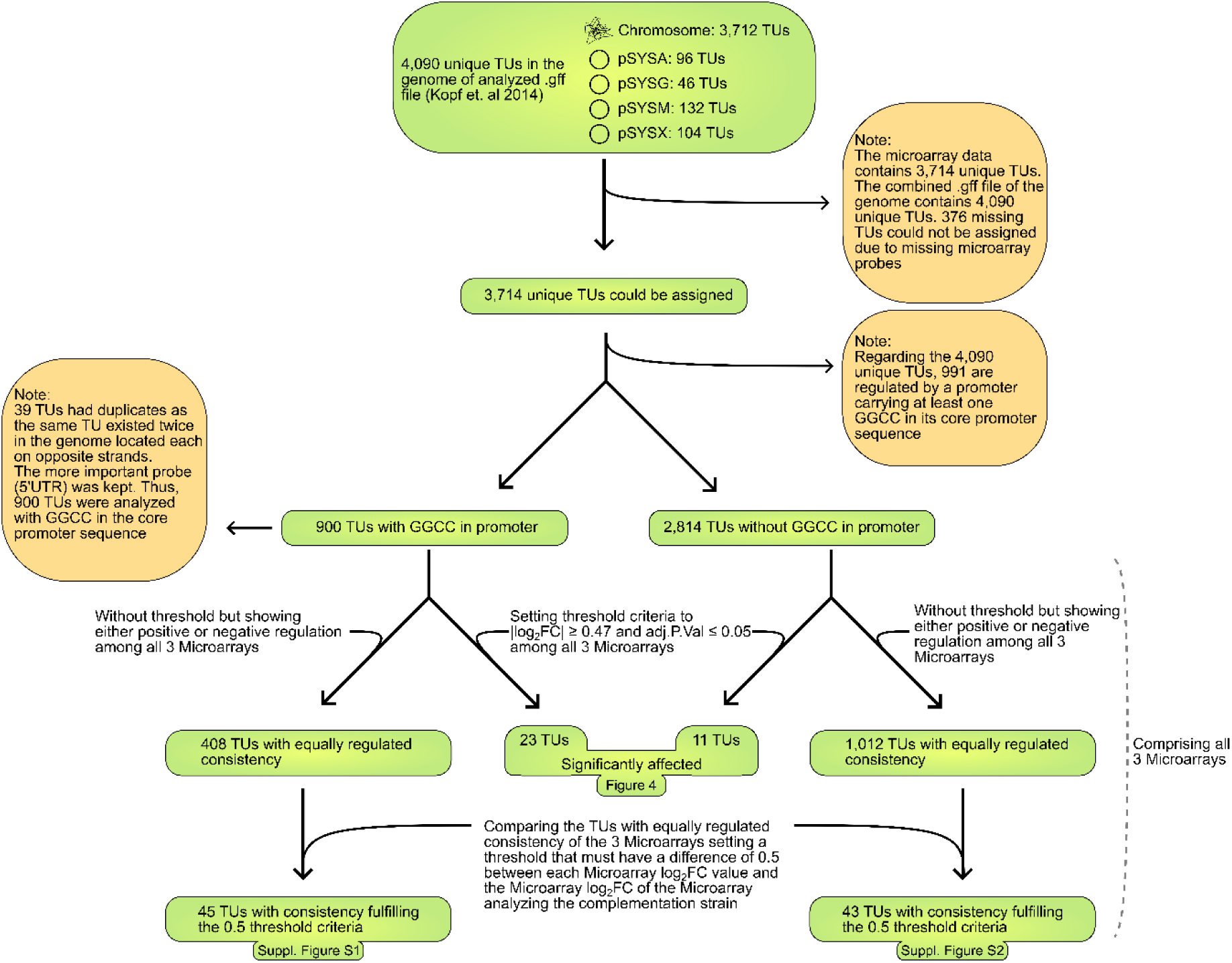
Work-flow of bioinformatic analyses. To analyze the occurrence of the GGCC motif within promoter sequences, the dataset of experimentally mapped TSSs including the chromosome and the 4 large plasmids, pSYSA, pSYSG, pSYSM, and pSYSX (Kopf *et al*., 2014) was used. A sequence window spanning −65 to +11 relative to the annotated TSS was defined. For the prediction of −10 promoter elements, the position-specific weight matrix (PSWM) defined by Mitschke *et al*. (2011) was used. The PSWM was used for scoring hexanucleotide motifs as possible −10 elements with a 5′ end −14 to −10 nt relative to the TSS. The −10 element with the highest score was used.

## Acknowledgments

We thank Ingeborg Scholz, University of Freiburg, and Klaudia Radloff-Michl, University of Rostock, for excellent technical assistance. This study is based on the doctoral thesis of Nils Stappert. The work was funded by the German Research Foundation (DFG) (grant HE 2544/12-2 to WRH and grant HA 2002/17-2 to MH).

## Ethics Statement

The authors have nothing to report.

## Copyright Statement

No copyright permissions were needed to be taken into account.

## Data Availability

The previously generated microarray data in the GEO database at https://www.ncbi.nlm.nih.gov/geo/query/acc.cgi?acc=GSE126282, accession number GSE126282 (BioProject PRJNA521475) was used.

## Author Contributions

WRH and MH designed the study. VR performed transcriptomic analyses. NSt and NSc performed all other experiments and evaluated data. NSt conducted the bioinformatics analyses. WRH and MH contributed to data analyses. NSt, WRH and MH drafted the manuscript. All authors read and approved the final manuscript.

## Supplementary Tables

**Table S1.** List of *Synechocystis* 6803 strains and corresponding transcriptomic datasets used in this study.

| Name/Dataset | Description of strains | AB resistance | References |
|---|---|---|---|
| 2014 Com | Complementation mutant in which the Δ*sll0729* deletion is complemented by *sll0729* located on the vector pVZ322, under the transcriptional control of the Zn<sup>2+</sup>-inducible promoter P<sub>*ziaA*</sub> | Amp and Cm | (Hagemann *et al*., 2018; Gärtner *et al*., 2019) |
| 2014 Ko | Δ*sll0729* deletion mutant | Km |  |
| 2024 LL | Δ*sll0729* knockout mutants generated in the year 2024 grown under low light (15 µmol m<sup>-2</sup> s<sup>-1</sup>, 30 °C, 2h). Carrying a single mutation in the promoter sequence of P<sub>*slr1790*</sub> changing the cytosine at position −4 (chromosomal position 256,605 RefSeq., NC_000911) in the discriminator sequence (TSS +1) to thymine leading from GGCC to GGTC | Km | (Schmidt *et al*., 2024) |
| 2024 HL | Δ*sll0729* knockout mutants generated in the year 2024 grown under high light (500 µmol m<sup>-2</sup> s<sup>-1</sup>, 30° C, 30 min). Carrying a single mutation in the promoter sequence of P<sub>*slr1790*</sub> changing the cytosine at position −4 (chromosomal position 256,605 RefSeq., NC_000911) in the discriminator sequence (TSS +1) to thymine leading from GGCC to GGTC | Km |  |
| Δ*sll0729*N | Δ*sll0729* knockout mutant generated in the year 2024 | Km | (Schmidt *et al*., 2024) |
| Suppr. Δ*sll0729* | Suppressor mutant of Δ*sll0729* carrying the GGCC to GGTC mutation in the discriminator sequence of P<sub>*slr1790*</sub> at position −4 (chromosomal position 256,605 RefSeq., NC_000911) in the discriminator sequence (TSS +1) | Km | (Gärtner *et al*., 2019; Schmidt *et al*., 2024) |

**Table S2.**
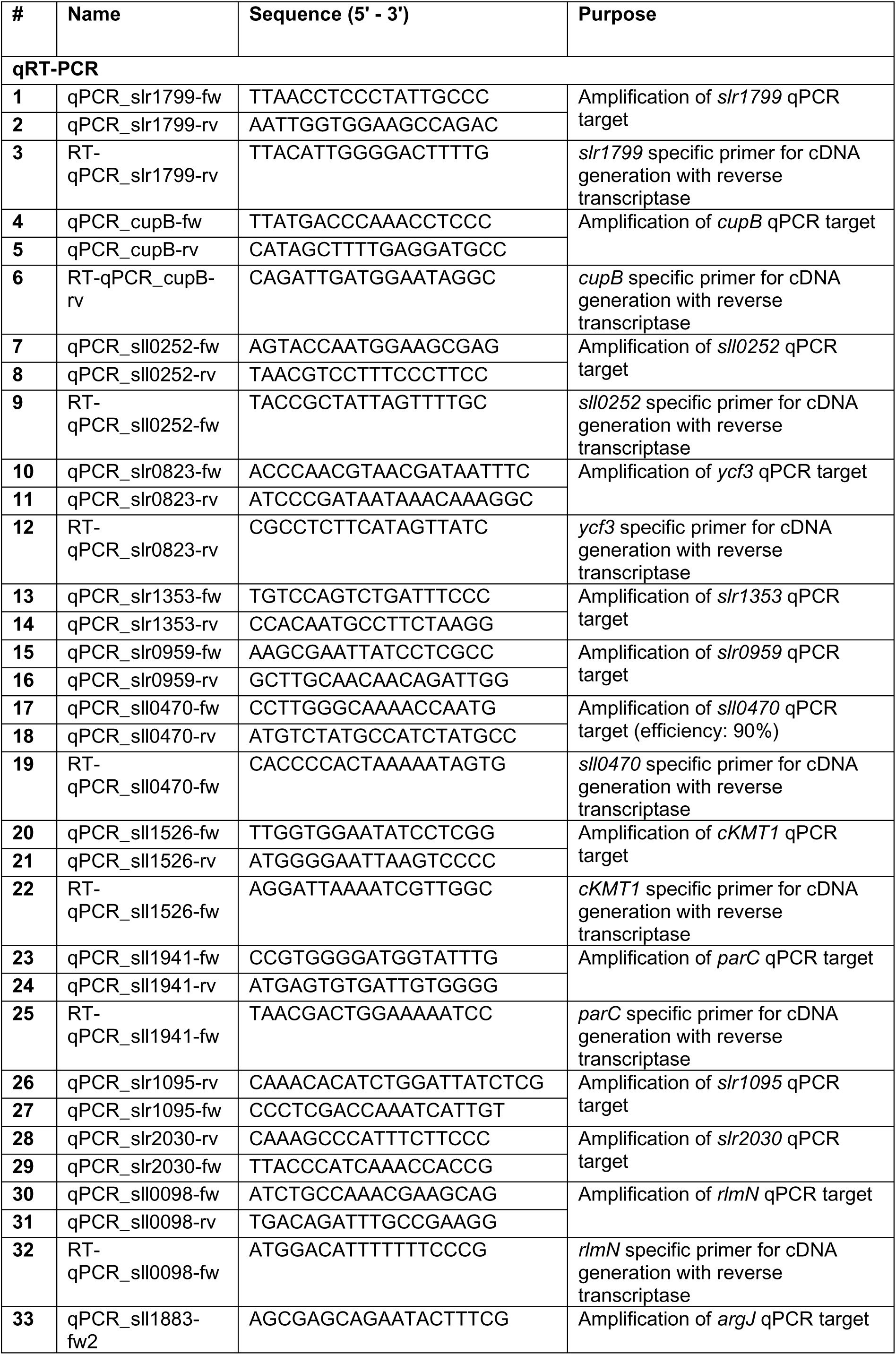
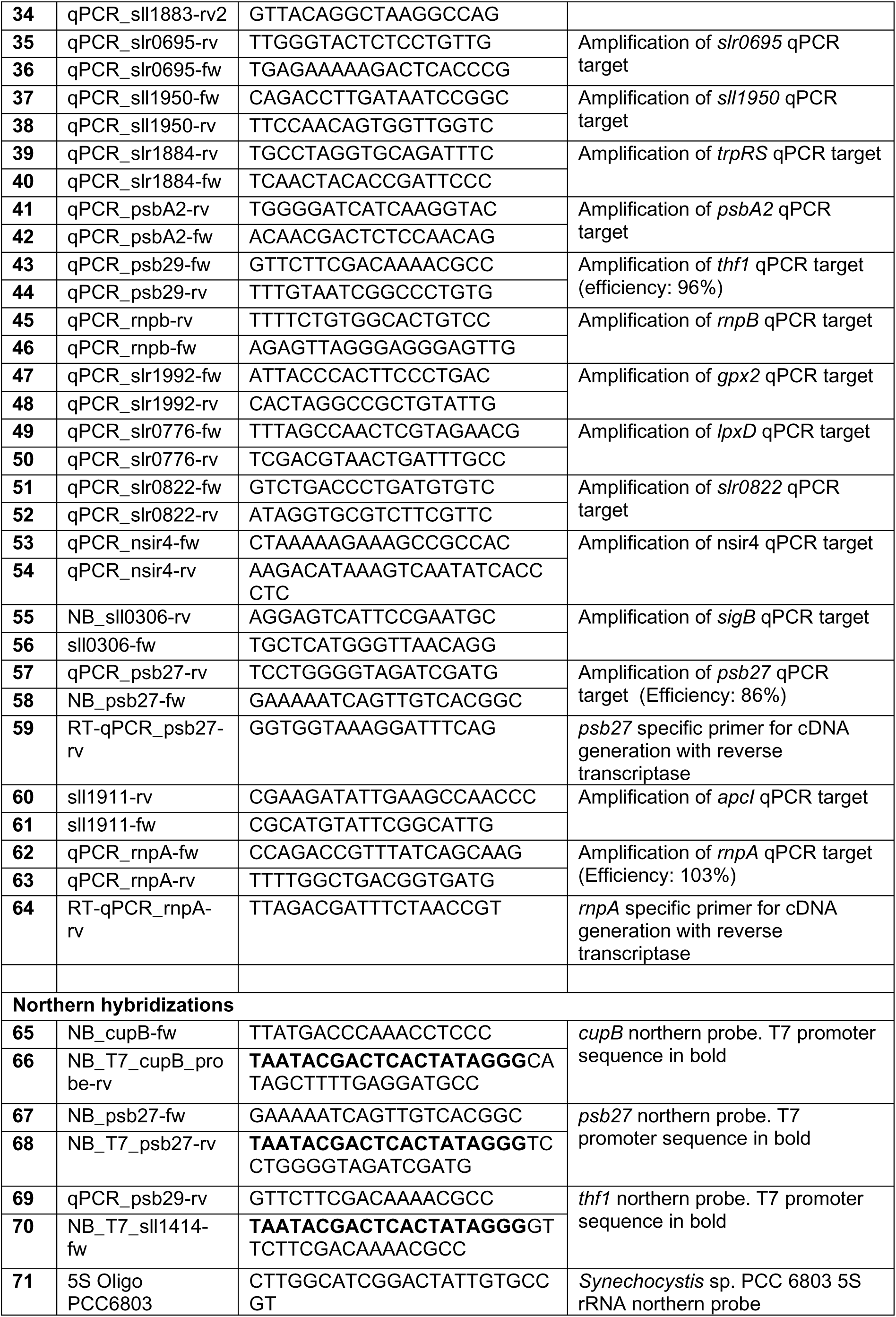
List of oligonucleotides used in this study.

## References

Alvarenga-Lucius, L., Maaß, S., Reimann, V., Becher, D., Hess, W.R., and Hagemann, M. (2026) Impact of SliP4 deletion on the high-light acclimation in *Synechocystis* sp. PCC 6803. Microlife 7: uqag003.

Asayama, M., and Imamura, S. (2008) Stringent promoter recognition and autoregulation by the group 3 σ-factor SigF in the cyanobacterium *Synechocystis* sp. strain PCC 6803. Nucleic Acids Research 36: 5297–5305.

Benjamini, Y., and Hochberg, Y. (1995) Controlling the false discovery rate: a practical and powerful approach to multiple testing. Journal of the Royal Statistical Society: Series B (Methodological) 57: 289–300.

Blow, M.J., Clark, T.A., Daum, C.G., Deutschbauer, A.M., Fomenkov, A., Fries, R., et al. (2016) The epigenomic landscape of prokaryotes. PLOS Genetics 12: e1005854.

Camacho, E.M., and Casadesús, J. (2005) Regulation of *traJ* transcription in the *Salmonella* virulence plasmid by strand-specific DNA adenine hemimethylation. Mol Microbiol 57: 1700–1718.

Cleveland, W.S., and Devlin, S.J. (1988) Locally weighted regression: an approach to regression analysis by local fitting. Journal of the American Statistical Association 83: 596–610.

Dhe-Paganon, S., Syeda, F., and Park, L. (2011) DNA methyl transferase 1: regulatory mechanisms and implications in health and disease. Int J Biochem Mol Biol 2: 58–66.

Elhai, J. (2015) Highly iterated palindromic sequences (HIPs) and their relationship to DNA methyltransferases. Life 5: 921–948.

Flombaum, P., Gallegos, J.L., Gordillo, R.A., Rincón, J., Zabala, L.L., Jiao, N., et al. (2013) Present and future global distributions of the marine cyanobacteria *Prochlorococcus* and *Synechococcus*. Proc Natl Acad Sci U S A 110: 9824–9829.

Gärtner, K., Klähn, S., Watanabe, S., Mikkat, S., Scholz, I., Hess, W.R., and Hagemann, M. (2019) Cytosine N4-methylation via M.Ssp6803II is involved in the regulation of transcription, fine-tuning of DNA replication and DNA repair in the cyanobacterium *Synechocystis* sp. PCC 6803. Frontiers in Microbiology 10: 1–14.

Gaultney, R.A., Vincent, A.T., Lorioux, C., Coppée, J.-Y., Sismeiro, O., Varet, H., et al. (2020) 4-Methylcytosine DNA modification is critical for global epigenetic regulation and virulence in the human pathogen *Leptospira interrogans*. Nucleic Acids Res 48: 12102– 12115.

Giner-Lamia, J., Robles-Rengel, R., Hernández-Prieto, M.A., Muro-Pastor, M.I., Florencio, F.J., and Futschik, M.E. (2017) Identification of the direct regulon of NtcA during early acclimation to nitrogen starvation in the cyanobacterium *Synechocystis* sp. PCC 6803. Nucleic Acids Research 45: 11800–11820.

Hagemann, M., Gärtner, K., Scharnagl, M., Bolay, P., Lott, S.C., Fuss, J., et al. (2018) Identification of the DNA methyltransferases establishing the methylome of the cyanobacterium *Synechocystis* sp. PCC 6803. DNA Res 25: 343–352.

Hagemann, M., and Hess, W.R. (2018) Systems and synthetic biology for the biotechnological application of cyanobacteria. Curr Opin Biotechnol 49: 94–99.

Hagemann, M., Song, S., and Brouwer, E. (2021) Inorganic carbon assimilation in cyanobacteria: mechanisms, regulation, and engineering. In Cyanobacteria Biotechnology. Nielsen, J., Lee, S., Stephanopoulos, G., and Hudson, P. (eds). Wiley, pp. 1–31 https://onlinelibrary.wiley.com/doi/10.1002/9783527824908.ch1. Accessed April 24, 2026.

Hihara, Y., Kamei, A., Kanehisa, M., Kaplan, A., and Ikeuchi, M. (2001) DNA microarray analysis of cyanobacterial gene expression during acclimation to high light. Plant Cell 13: 793–806.

Hu, L., Xiao, P., Jiang, Y., Dong, M., Chen, Z., Li, H., et al. (2018) Transgenerational epigenetic inheritance under environmental stress by genome-wide DNA methylation profiling in cyanobacterium. Front Microbiol 9: 1479.

Imamura, S., and Asayama, M. (2009) Sigma factors for cyanobacterial transcription. Gene Regul Syst Bio 3: 65–87.

Iyer, L.M., Zhang, D., and Aravind, L. (2016) Adenine methylation in eukaryotes: Apprehending the complex evolutionary history and functional potential of an epigenetic modification. BioEssays 38: 27–40.

Jeltsch, A. (2002) Beyond Watson and Crick: DNA methylation and molecular enzymology of DNA methyltransferases. Chembiochem 3: 274–293.

Jin, B., and Robertson, K.D. (2013) DNA methyltransferases, DNA damage repair, and cancer. Adv Exp Med Biol 754: 3–29.

Kahramanoglou, C., Prieto, A.I., Khedkar, S., Haase, B., Gupta, A., Benes, V., et al. (2012) Genomics of DNA cytosine methylation in *Escherichia coli* reveals its role in stationary phase transcription. Nat Commun 3: 886.

Klähn, S., Opel, F., and Hess, W.R. (2024) Customized molecular tools to strengthen metabolic engineering of cyanobacteria. Green Carbon 2: 149–163.

Klähn, S., Schaal, C., Georg, J., Baumgartner, D., Knippen, G., Hagemann, M., et al. (2015) The sRNA NsiR4 is involved in nitrogen assimilation control in cyanobacteria by targeting glutamine synthetase inactivating factor IF7. Proc Natl Acad Sci USA 112 https://pnas.org/doi/full/10.1073/pnas.1508412112. Accessed May 7, 2025.

Konert, M.M., Wysocka, A., Koník, P., and Sobotka, R. (2022) High-light-inducible proteins HliA and HliB: pigment binding and protein-protein interactions. Photosynth Res 152: 317–332.

Kopf, M., Klähn, S., Scholz, I., Matthiessen, J.K.F., Hess, W.R., and Voß, B. (2014) Comparative analysis of the primary transcriptome of *Synechocystis* sp. PCC 6803. DNA Res 21: 527–539.

Militello, K.T., Simon, R.D., Qureshi, M., Maines, R., VanHorne, M.L., Hennick, S.M., et al. (2012) Conservation of Dcm-mediated cytosine DNA methylation in *Escherichia coli*. FEMS Microbiol Lett 328: 78–85.

Mitschke, J., Georg, J., Scholz, I., Sharma, C.M., Dienst, D., Bantscheff, J., et al. (2011) An experimentally anchored map of transcriptional start sites in the model cyanobacterium *Synechocystis* sp. PCC6803. Proc Natl Acad Sci USA 108: 2124–2129.

Moore, L.D., Le, T., and Fan, G. (2013) DNA methylation and its basic function. Neuropsychopharmacol 38: 23–38.

Murphy, J., Mahony, J., Ainsworth, S., Nauta, A., and Sinderen, D. van (2013) Bacteriophage orphan DNA methyltransferases: insights from their bacterial origin, function, and occurrence. Appl Environ Microbiol 79: 7547–7555.

Nikkanen, L., Vakal, S., Hubáček, M., Santana-Sánchez, A., Konert, G., Wang, Y., et al. (2025) Flavodiiron proteins associate pH-dependently with the thylakoid membrane for ferredoxin-1-powered O2 photoreduction. New Phytol 246: 2084–2101.

Pinto, F.L., Thapper, A., Sontheim, W., and Lindblad, P. (2009) Analysis of current and alternative phenol based RNA extraction methodologies for cyanobacteria. BMC Molecular Biol 10: 79.

Rippka, R., Deruelles, J., Waterbury, J.B., Herdman, M., and Stanier, R.Y. (1979) Generic assignments, strain histories and properties of pure cultures of cyanobacteria. Microbiology 111: 1–61.

Ritchie, M.E., Phipson, B., Wu, D., Hu, Y., Law, C.W., Shi, W., and Smyth, G.K. (2015) limma powers differential expression analyses for RNA-sequencing and microarray studies. Nucleic Acids Research 43: e47–e47.

Roberts, R.J., Vincze, T., Posfai, J., and Macelis, D. (2010) REBASE—a database for DNA restriction and modification: enzymes, genes and genomes. Nucleic Acids Research 38: D234–D236.

Rodriguez, F., Yushenova, I.A., DiCorpo, D., and Arkhipova, I.R. (2022) Bacterial N4-methylcytosine as an epigenetic mark in eukaryotic DNA. Nat Commun 13: 1072.

Sánchez-Romero, M.A., and Casadesús, J. (2020) The bacterial epigenome. Nat Rev Microbiol 18: 7–20.

Schmidt, N., Stappert, N., Nimura-Matsune, K., Watanabe, S., Sobotka, R., Hagemann, M., and Hess, W.R. (2024) Epigenetic control of tetrapyrrole biosynthesis by m4C DNA methylation in a cyanobacterium. DNA Res 31: dsae035.

Scholz, I., Lange, S.J., Hein, S., Hess, W.R., and Backofen, R. (2013) CRISPR-Cas systems in the cyanobacterium *Synechocystis* sp. PCC6803 exhibit distinct processing pathways involving at least two Cas6 and a Cmr2 protein. PLoS ONE 8: e56470.

Sergeeva, A., Davydova, K., Perenkov, A., and Vedunova, M. (2023) Mechanisms of human DNA methylation, alteration of methylation patterns in physiological processes and oncology. Gene 875: 147487.

Srivastava, A., Summers, M.L., and Sobotka, R. (2020) Cyanobacterial sigma factors: Current and future applications for biotechnological advances. Biotechnology Advances 40: 107517.

Trautmann, D., Voß, B., Wilde, A., Al-Babili, S., and Hess, W.R. (2012) Microevolution in cyanobacteria: re-sequencing a motile substrain of *Synechocystis* sp. PCC 6803. DNA Res 19: 435–448.

Voß, B., and Hess, W.R. (2014) The Identification of Bacterial Non-coding RNAs through Complementary Approaches. In Handbook of RNA Biochemistry. Hartmann, R.K., Bindereif, A., Schön, A., and Westhof, E. (eds). Wiley, pp. 787–800 https://onlinelibrary.wiley.com/doi/10.1002/9783527647064.ch34. Accessed February 7, 2026.

Woude, M. van der, Braaten, B., and Low, D. (1996) Epigenetic phase variation of the *pap* operon in *Escherichia coli*. Trends Microbiol 4: 5–9.

